# Mechanistic studies of non-canonical amino acid mutagenesis

**DOI:** 10.1101/2021.05.24.445427

**Authors:** Rachel C. Fleisher, Nina Michael, Ruben L. Gonzalez

## Abstract

Over the past decade, harnessing the cellular protein synthesis machinery to incorporate non-canonical amino acids (ncAAs) into tailor-made peptides has significantly advanced many aspects of molecular science. More recently, groundbreaking progress in our ability to engineer this machinery for improved ncAA incorporation has led to significant enhancements of this powerful tool for biology and chemistry. By revealing the molecular basis for the poor or improved incorporation of ncAAs, mechanistic studies of ncAA incorporation by the protein synthesis machinery have tremendous potential for informing and directing such engineering efforts. In this chapter, we describe a set of complementary biochemical and single-molecule fluorescence assays that we have adapted for mechanistic studies of ncAA incorporation. Collectively, these assays provide data that can guide engineering of the protein synthesis machinery to expand the range of ncAAs that can be incorporated into peptides and increase the efficiency with which they can be incorporated, thereby enabling the full potential of ncAA mutagenesis technology to be realized.

## 1. Introduction

The field of non-canonical amino acid (ncAA) mutagenesis aims to harness the power of the cellular machinery that translates messenger RNAs (mRNAs) into proteins, which we refer to here as the ‘translation machinery’ (TM), to make templated designer peptides with amino acids other than the twenty canonical amino acids (cAAs) (Liu & Schultz, 2010). The applications of this technology are diverse, including the creation of difficult-to-synthesize therapeutics (Obexer et al., 2017); novel materials (Katoh et al., 2020; Rogers et al., 2018); and proteins engineered to contain biophysical probes (Braun et al., 2019; Chung et al., 2020; Desai & Gonzalez, 2020; Lee et al., 2019; Saleh et al., 2019) or perform new functions (Drienovská & Roelfes, 2020; Wang, 2017). Despite this, many classes of ncAAs are poor substrates for the TM (Pavlov et al., 2009; Tan et al., 2004) and improving their incorporation into peptides often requires engineering of the TM (Amiram et al., 2015; Dedkova & Hecht, 2019; Fan et al., 2015; Hammerling et al., 2020; Takayuki Katoh et al., 2017; Lajoie et al., 2013; Ohtsuki et al., 2010; Park et al., 2011; Rackham & Chin, 2005; Rogers et al., 2018; Tharp et al., 2020; Tsiamantas et al., 2020). Recent breakthroughs in the synthesis of ncAA-transfer RNAs (tRNAs) (Bianco et al., 2012; Desai & Gonzalez, 2020; Dunkelmann et al., 2020; Ernst et al., 2016; Liu & Schultz, 2010; Liu et al., 2017; Murakami et al., 2006; Neumann et al., 2010; Wang et al., 2001) and the engineering of tRNAs (Katoh & Suga, 2018; T. Katoh et al., 2017), translation factors (Katoh & Suga, 2018; T. Katoh et al., 2017; Park et al., 2011), and the ribosome itself (Aleksashin et al., 2020; Chen et al., 2017; Dedkova et al., 2003, 2006; Dedkova et al., 2012; Fried et al., 2015; Liu et al., 2014; Maini, Dedkova, et al., 2015; Neumann et al., 2010; Orelle et al., 2015; Schmied et al., 2018), particularly in the context of cell strains optimized for ncAA incorporation (Isaacs et al., 2011; Lajoie et al., 2013), have pushed the field to new heights. These developments have facilitated the incorporation of several classes of ncAAs that had previously been poorly tolerated by the TM, increased the diversity of ncAAs that can be used by the TM, and enabled the incorporation of multiple ncAAs into single peptides, both *in vitro* and *in vivo*, in bacterial as well as eukaryotic cells.

Despite the success of these engineering efforts, the mechanistic basis through which they improve ncAA incorporation is not always understood, limiting our ability to improve and expand them. Specifically, determining when, where, and how during translation the TM fails to incorporate particular ncAAs can inform rational design-or directed evolution approaches aimed at overcoming such mechanistic obstacles and expanding the range of ncAAs that the TM can use. Motivated by this principle, we and others are using biochemistry (Aleksashin et al., 2019; Effraim et al., 2009; Englander et al., 2015; Fleisher et al., 2018; Gamper et al., 2021; Heckler et al., 1988), rapid kinetics (Gamper et al., 2021; Liljeruhm et al., 2019), single-molecule biophysics (Effraim et al., 2009; Gamper et al., 2021), structural biology (Melnikov et al., 2019; Schmied et al., 2018; Ward et al., 2019), molecular dynamic (MD) simulations (Englander et al., 2015; Maini, Chowdhury, et al., 2015), computational analyses (Walker et al., 2020), and other mechanistic tools to explore the limits of ncAA incorporation by the TM.

As an example from our own work, we have found that once a D-amino acid (aa) is incorporated into the C-terminal end of the nascent polypeptide chain being synthesized by the bacterial TM, the incorporated D-aa arrests translation by perturbing the conformational dynamics of the ribosomal peptidyl transferase center (PTC) (Englander et al., 2015), consistent with, and extending, earlier findings by Hecht and co-workers (Heckler et al., 1988). Remarkably, the ability of the D-aa to perturb these dynamics can be modulated by both the tRNA- and amino acid components of the next, incoming aa-tRNA (Fleisher et al., 2018). The findings suggested that engineering specific features of the PTC and/or incoming aa-tRNA can facilitate incorporation of D-aas and, very likely, other ncAAs that might arrest translation through a similar mechanism. In a second example, we have investigated the mechanism through which a tRNA containing a one-nucleotide insertion mutation in its anticodon loop induces the bacterial TM to undergo a highly efficient +1 frameshifting (FS) event at a specific quadruplet-nucleotide codon, thereby enabling the robust incorporation of an ncAA by the TM (Gamper et al., 2021). In this study, we found that the tRNA induces +1FS at the quadruplet codon by manipulating a specific conformational rearrangement of the ribosomal small, or 30S, subunit that is necessary for translocation of the ribosome along the mRNA. Our findings therefore provide a guide for engineering specific structural elements of the 30S subunit so as to facilitate the +1FS tRNA-mediated incorporation of ncAAs in response to quadruplet codons.

We begin this chapter with a short description of the general approaches we and our collaborators have used to prepare ncAA-tRNAs for our studies, providing references to appropriately detailed protocols for the preparation of these essential reagents. We next briefly describe the reconstituted *Escherichia coli in vitro* translation system we have reported in a previous *Methods in Enzymology* chapter (Fei et al., 2010) and have used in our studies. The bulk of the chapter then provides detailed descriptions of the set of biochemistry and single-molecule fluorescence resonance energy transfer (smFRET) assays we have used in our mechanistic studies of ncAA incorporation by the TM. We close by highlighting a number of mechanistic tools that are only just now beginning to be applied in earnest to studies of ncAA incorporation, but that we think hold tremendous promise for ongoing and future efforts to engineer the TM for improved ncAA mutagenesis capabilities.

## 2. Preparation of ncAA-tRNAs

### 2.1. Overview of methods used to prepare ncAA-tRNAs

Several methods exist for aminoacylating tRNAs with ncAAs. A common method for preparation of ncAA-tRNAs involves a hybrid strategy of chemically synthesizing the 5’-phospho-2’-deoxycytidylyl-(3’,5’)-adenosine (pdCpA) dinucleotide and acylating it with the cyanomethyl active ester (CME) of an α-amine-protected amino acid. T4 RNA ligase is then used to ligate the resulting pdCpA-aa to the 3’ end of an *in vitro* transcribed tRNA lacking the universally conserved cytidine (C) and adenine (A) nucleotides at positions 75 and 76 at its 3’ end as described by Robertson et al. (1991). tRNA synthetases (aaRS) that have been engineered so as to expand their substrate specificities are also commonly used for the preparation of ncAA-tRNAs (Datta et al., 2002; Melnikov & Söll, 2019). Because of its ease of use and the fact that it allows us to use naturally occurring tRNAs containing post-transcriptional modifications that reduce unwanted, tRNA-induced frameshifting (Hou et al., 2015), we typically employ the ribozyme-based ‘Flexizyme’ system developed by Suga and co-workers (Murakami et al., 2006; Ohuchi et al., 2007) to prepare our ncAA-tRNAs.

### 2.2. Reagents, equipment, and general procedures for preparing ncAA-tRNAs using the Flexizyme system

The following reagents and equipment were used for preparing ncAA-tRNAs using the Flexizyme system. Chemical reagents, enzymes, and tRNAs were purchased from Sigma Aldrich unless otherwise specified. Analytical high-performance liquid chromatography (HPLC) was executed using a reversed-phased Phenomenex Kinetex C18 column on a Waters 600 HPLC system or a reversed-phase Waters Xbridge C18 column on a Shimadzu LC-10AD^VP^ HPLC system. Preparative HPLC was performed using a reversed-phase Phenomenex Luna C18(2) column on a Waters 600 HPLC system. Proton (^1^H) nuclear magnetic resonance (NMR) spectroscopy was executed on a Bruker DPX-400 instrument. Analogous HPLC and NMR instrumentation may be used in place of the ones listed above. In addition, if necessary, appropriate substitutions may be made for some of the equipment and instruments listed below.

#### Chemical reagents, solvents, and buffer salts

α-N-tert-butyloxycarbonyl (α-N-Boc)-protected amino acids (from Chem Impex)

1-fluoro-2-4-dinitrophenyl-5-L-alanine amide (FDAA, Marfey’s Reagent)

1,8-bis(dimethylamino)naphthalene (Proton Sponge)

3,5-Dinitrobenzyl chloride

Chloroacetonitrile (ClCH_2_CN)

Triethylamine (TEA)

Dimethylformamide (DMF)

Diethyl ether (Et_2_O)

Hydrochloric acid (HCl)

Sodium bicarbonate (NaHCO_3_)

Saturated sodium chloride (NaCl) solution (brine)

Magnesium sulfate (MgSO_4_)

Ammonium acetate (NH_4_OAc)

Methanol (MeOH)

Ethanol (EtOH)

Ethyl acetate (EtOAc)

Acetonitrile (MeCN)

Trifluoroacetic acid (TFA)

Potassium 2-[4-(2-hydroxyethyl)piperazin-1-yl]ethanesulfonic acid (K-HEPES)

Potassium chloride (KCl)

Magnesium chloride (MgCl_2_)

Dimethyl sulfoxide (DMSO)

Potassium acetate (KOAc)

Tris hydrochloride (Tris-HCl; pH_RT_ = 8.0)

Ammonium chloride (NH_4_Cl)

Acetic acid (HOAc)

[^32^P]adenosine-5’-monophosphate (Perkin Elmer)

#### Ribozymes, tRNAs, and enzymes

Dinitro-Flexizyme (dFx) (prepared as described in Murakami et al. (2006))

Enhanced Flexizyme (eFx) (prepared as described in Murakami et al. (2006)) aa-specific tRNAs

*E. coli* nucleotidyl transferase (overexpressed from a plasmid kindly provided by Dr. Ya-Ming Hou (Thomas Jefferson University) and prepared as described in Dupasquier et al. (2008))

*Penicillium citrinum* nuclease P1

#### Equipment and instruments

Vacuum gas manifold

High vacuum pump

Büchner funnel and filter flask

Phosphorimaging screen (GE Healthcare)

Phosphorimager (Typhoon FLA7000; GE Healthcare)

#### Other supplies

Molecular sieves (3Å)

Polyethyleneimine-impregnated (PEI)-cellulose thin layer chromatography (TLC) plates (EMD Chemicals)

Saran wrap

Image analysis software (ImageQuant, ImageJ, or similar)

### 2.3. Preparation of ncAA-tRNAs using the Flexizyme system

#### 2.3.1. Overview of amino acid active ester preparation

Starting from commercially sourced α-N-Boc-protected amino acids, we follow the protocols previously published by Suga and coworkers (Murakami et al., 2006) to synthesize the amino acid 3,5-dinitrobenyl esters (DBEs) and CMEs we use with the Flexizyme system (Avins, 2010; Effraim et al., 2009; Englander et al., 2015; Fleisher et al., 2018). The only exception to this was for our synthesis of D-phenylalanine (Phe)-CME, in which we modified the protocol in Murakami et al. (2006) to limit the extent of racemization (*vide infra*) (Avins, 2010). For the syntheses of D-aa active esters, we verified the enantiomeric excess for each D-aa active ester to be above 98%, as determined by Marfey’s analysis (Adamson et al., 1992; Kochhar et al., 2000), a well-established technique for assessing the stereochemical purity of amino acids and peptides (Goodlett et al., 1995). Reaction with the chiral Marfey’s Reagent converts the enantiomeric D-aa active ester to a diastereomer that is easily separable using reversed-phase C18 column chromatography in a standard HPLC system outfitted with an ultraviolet (UV) detector set to detect the FDAA moiety at 340 nm. We found that quantifying the stereochemical purity of the D-aa active esters was critically important because L-aa-tRNAs are incorporated by the TM with much greater efficiency than their D-aa-tRNA counterparts and contaminating L-aa-tRNA can thereby lead to misinterpretation of the results. We further characterized the products of our peptide synthesis assays (*vide infra*) by HPLC, using comigration of our products with chemically synthesized, D-aa-containing, ‘authentic’, marker peptides to ensure that no racemization took place during aminoacylation with the Flexizyme. The excellent agreement between the optical purity of the D-aa active esters and the stereochemical assessment of the peptide products was consistent with no further racemization taking place in the steps following the D-aa active ester synthesis (Avins, 2010).

#### 2.3.2. Representative protocol for syntheses of ncAA-DBEs (D-Lys-DBE)

A mixture of α-N-Boc-D-Lysine (Lys) (300 mg, 1.6 mmol), 3,5-dinitrobenzyl chloride (286 mg, 1.3 mmol), and TEA (270 mg, 2.7 mmol) in 2.0 mL of DMF is allowed to react at room temperature overnight. The reaction is then diluted with Et_2_O (30 mL) and the solution extracted with 0.5 M HCl (10 mL x 3), 4% NaHCO_3_ (10 mL x 3), and brine (20 mL x1). The organic layer is isolated, dried with MgSO_4_ and concentrated. To remove the Boc protecting group, the resulting crude residue is then dissolved in 8 mL of 4 N HCl in EtOAc and stirred for 20 min at room temperature. The solution is concentrated again and any remaining HCl is removed by high vacuum. The product is precipitated by the addition of a Et_2_O:MeOH mixture (10:1 v/v) and the precipitate is filtered (Avins, 2010; Murakami et al., 2006). In our hands, we achieve a 30% overall yield (Avins, 2010). ^1^H NMR (400 MHz, CD_3_OD): δ 9.03 (t, 1H, Ar-C_4_H), 8.73 (d, 2H, Ar-C_2,6_H), 5.56 (s, 2H, OCH_2_), 4.24 (t, 1H_α_), 2.97 (t, 2H, N_ε_CH_2_), 2.13-1.94 (m, 2H), 1.75 (q, 2H), 1.68-1.51 (m, 2H) (Avins, 2010).

#### 2.3.3. Synthesis of D-Phe-CME

A mixture of α-N-Boc-D-Phe (265 mg, 1 mmol), Proton Sponge (429 mg, 2 mmol), and ClCH_2_CN (1 mL, excess, dried over molecular sieves) is stirred under inert atmosphere at room temperature overnight. EtOAc (36 mL) is added to the reaction. The resulting solution is extracted and the Boc protecting group is removed as described for the DBE syntheses in Section 2.3.2. The product is obtained in 35% overall yield (125 mg, 0.47 mmol) and is then further purified by semi-preparative HPLC using a reversed-phase C18 column and a gradient of 1% MeCN and 0.1% TFA to 80% MeCN and 0.1% TFA over 75 min ((Avins, 2010); modified from Murakami et al. (2006)). ^1^H NMR (400 MHz, CD_3_OD): δ 7.44 (m, 3H, Ar-C_3,4,5_H), 7.32 (d, 2H, Ar-C_2,6_H), 5.05 (s, 2H, OCH_2_CN), 3.3 (m, 2H, CH_2_-Ar) (Avins, 2010).

#### 2.3.4. Aminoacylation using Flexizymes

tRNAs are aminoacylated using amino acid DBEs or CMEs and dFx or eFx, respectively (Effraim et al., 2009; Englander et al., 2015; Fleisher et al., 2018; Murakami et al., 2006). The aminoacylation reactions contain 20 μM tRNA, 20 μM dFx or eFx, and 5 mM amino acid DBEs or CMEs, respectively, in a buffer of 0.1 M K-HEPES (pH_RT_ = 7.5), 0.1 M KCl, 600 mM MgCl_2_, and 20% DMSO. Reactions proceed on ice, with the length of time determined by the identity of the amino acid side chain. The reactions are subsequently quenched with 3 volumes of 600 mM NH_4_OAc (pH_RT_ = 5.0) and the ncAA-tRNA products are precipitated with EtOH. The ncAA-tRNAs in the resulting pellet are then resuspended and stored in 10 mM KOAc (pH_RT_ = 5.0) at –80 °C, and used without further purification. We noted that valine (Val)-specific tRNA (tRNA^Val^) purchased from Sigma was acylated to a certain extent with Val and therefore had to be deacylated prior to aminoacylation. Thus, following an earlier protocol reported by Powers and Noller (1991), tRNA^Val^ purchased from Sigma was treated with 1.8 M Tris-HCl (pH_RT_ = 8.0) for 3 h at 37 °C prior to using it in aminoacylation reactions.

To assess aminoacylation efficiency, analytical-scale aminoacylation reactions (5 μL total reaction volume) using 3’-[^32^P]-labeled tRNA are executed side-by-side with the preparative-scale reaction under identical conditions. The 3’ end of tRNA is [^32^P]-labeled with [^32^P]AMP using nucleotidyl transferase as described previously (Effraim et al., 2009; Ledoux & Uhlenbeck, 2008). After precipitation and resuspension, the reaction is digested with nuclease P1 for 10 min at room temperature. Separation of [^32^P]AMP from aa-[^32^P]AMP is achieved by TLC on PEI-cellulose TLC plates that have been pre-rinsed with water and dried, using a running buffer of 100 mM NH_4_Cl and 10% HOAc. TLC plates are then air-dried on a flat surface, wrapped in Saran wrap, exposed to a phosphorimaging screen overnight, and imaged using a phosphorimager (Effraim, 2010). Subsequently, the intensities of TLC spots corresponding to the unreacted [^32^P]AMP (*I*_AMP_) and aminoacylated product (*I*_aa-AMP_) are quantified using appropriate image analysis software. The aminoacylation efficiencies are calculated as [(*I*_aa-AMP_) / (*I*_AMP_ + *I*_aa-AMP_)] × 100. Experiments are typically executed in duplicate or triplicate, and the mean aminoacylation efficiency is reported along with the standard error of the mean in the case of duplicate measurements or the standard deviation in the case of triplicate measurements.

## 3. Assessing the performance of ncAA-tRNAs in translation

### 3.1. Preparation of a reconstituted, *E. coli in vitro* translation system

For our biochemical studies of translation using ncAA-tRNA substrates (Effraim et al., 2009; Englander et al., 2015; Fleisher et al., 2018), we use an *in vitro* translation system reconstituted from highly purified, *E. coli* components that, unless otherwise specified here, are prepared following detailed protocols we have published in a previous *Methods in Enzymology* chapter (Fei et al., 2010) that draws from Fei (2010). Briefly, tight-coupled 70S ribosomes, 30S subunits, and/or ribosomal large, or 50S, subunits are purified from *E. coli* MRE600 cells. tRNAs are purchased from Sigma, MP Biomedicals, or Chemical Block or, if commercially unavailable, are purified following straightforward adaptations (Effraim et al., 2009) of previously published protocols ((Louie et al., 1984; Ribeiro et al., 1995) or, more recently, (Kazayama et al., 2015; Tsurui et al., 1994)). Methionyl (Met)-tRNA synthetase (MetRS), PheRS, LysRS, and AlaRS, as well as Met-tRNA^fMet^ formyltransferase (FMT) are recombinantly expressed and purified from *E. coli* BL21(DE3) cells. Note that recombinant expression and purification of AlaRS is reported in (Effraim et al., 2009) rather than (Fei et al., 2010) and that additional aaRSs can be recombinantly expressed and purified using straightforward extensions of the protocols reported in Fei et al. (2010). Purchased or purified tRNAs and purified aaRSs and FMT are used to prepare cAA-tRNAs, including formylmethionyl (fMet)-tRNA^fMet^, as detailed in Fei et al. (2010). ncAA-tRNAs, on the other hand, are prepared as described in Section 2.3.4. The efficiency of aminoacylation and, in the case of fMet-tRNA^fMet^, formylation reactions can be determined using hydrophobic interaction chromatography (HIC), as detailed in Fei et al. (2010); acidic polyacrylamide gel electrophoresis (PAGE), as detailed in Effraim et al. (2009); or TLC, as described in Section 2.3.4. mRNAs are based on the gene for bacteriophage T4 gene product 32 and encode either the first 224, 20, or 9 amino acids of the gene (T4gp32_1–224_, T4gp32_1–20_, or T4gp32_1–9_ mRNAs, respectively) and are prepared either by run-off, *in vitro* transcription of plasmids using T7 RNA polymerase (T4gp32_1–224_ and T4gp32_1–20_ mRNAs) or by chemical synthesis (T4gp32_1–9_ mRNA; IDT, Inc.). A detailed description and sequence information for the T4gp32_1–224_ mRNA, from which the T4gp32_1–20_ and T4gp32_1–9_ mRNAs are further derived, can be found in Blanchard (2002). Additional descriptions and sequence information for the T4gp32_1–20_ mRNA can be found in Fei (2010) and Englander et al. (2015) and, for the T4gp32_1–9_ mRNA, in Fei (2010) and Fei et al. (2008). Also, a general protocol for the preparation of these mRNAs can be found in Fei et al. (2010) and examples of specific protocols for the preparation of mRNAs used in our studies of ncAA incorporation by the TM can be found in Effraim et al. (2009), Englander et al. (2015), and Fleisher et al. (2018). Like the aaRSs and FMT, the translation factors initiation factor (IF) 1, IF2, and IF3 and elongation factor (EF)-Tu, EF-Ts, and EF-G are all recombinantly expressed and purified from *E. coli* BL21(DE3) cells.

As a buffer for our *in vitro* translation system, we use a variant of the *in vitro* translation buffer established by Kurland, Ehrenberg, and coworkers (Jelenc & Kurland, 1979; Pavlov & Ehrenberg, 1996; Wagner et al., 1982) that we call Tris-Polymix Buffer and that is composed of 50 mM Tris-acetate (Tris-OAc; pH_RT_ = 7.5), 100 mM KCl, 5 mM NH_4_OAc, 0.5 mM calcium acetate (Ca(OAc)_2_), 6 mM β-mercaptoethanol (BME), 5 mM putrescine-HCl, 1 mM spermidine free base, and is adjusted to 0–15 mM magnesium acetate (Mg(OAc)_2_), as specified (Fei et al., 2010).

### 3.2. Overview of the translation elongation cycle

In our mechanistic studies of ncAA incorporation by the TM, we are interested in following the path of the ncAA throughout the entire translation elongation cycle, which is composed of three major steps: aa-tRNA selection, peptidyl transfer, and translocation (Figure 1). During the aa-tRNA selection step, a ternary complex (TC) of the translational guanine nucleotide triphosphatase (trGTPase) EF-Tu, GTP, and aa-tRNA is delivered to the ribosomal aa-tRNA binding (A) site. Transfer of the nascent polypeptide chain from the peptidyl-tRNA at the ribosomal peptidyl-tRNA binding (P) site to the aa-tRNA at the A site during the peptidyl transfer step then generates a newly deacylated tRNA at the P site and a newly formed peptidyl-tRNA at the A site. The trGTPase EF-G then catalyzes translocation of the deacylated tRNA and peptidyl-tRNA at the P and A sites into the ribosomal tRNA exit (E) and P sites, respectively, advancing the mRNA along with the tRNAs so as to bring the next codon into the now otherwise empty A site and ultimately prompting the release of the deacylated tRNA from the E site. Note that two rounds of the elongation cycle are required to fully test a single aa’s performance in all steps of the elongation cycle; its abilities to act as an acceptor or donor in the peptidyl transfer reaction are tested in the first and second cycles, respectively. In the sections that follow below, we will describe a battery of biochemical and smFRET assays that allow us to investigate the sub-steps of the elongation cycle in exceptional mechanistic detail.

**Figure 1.**
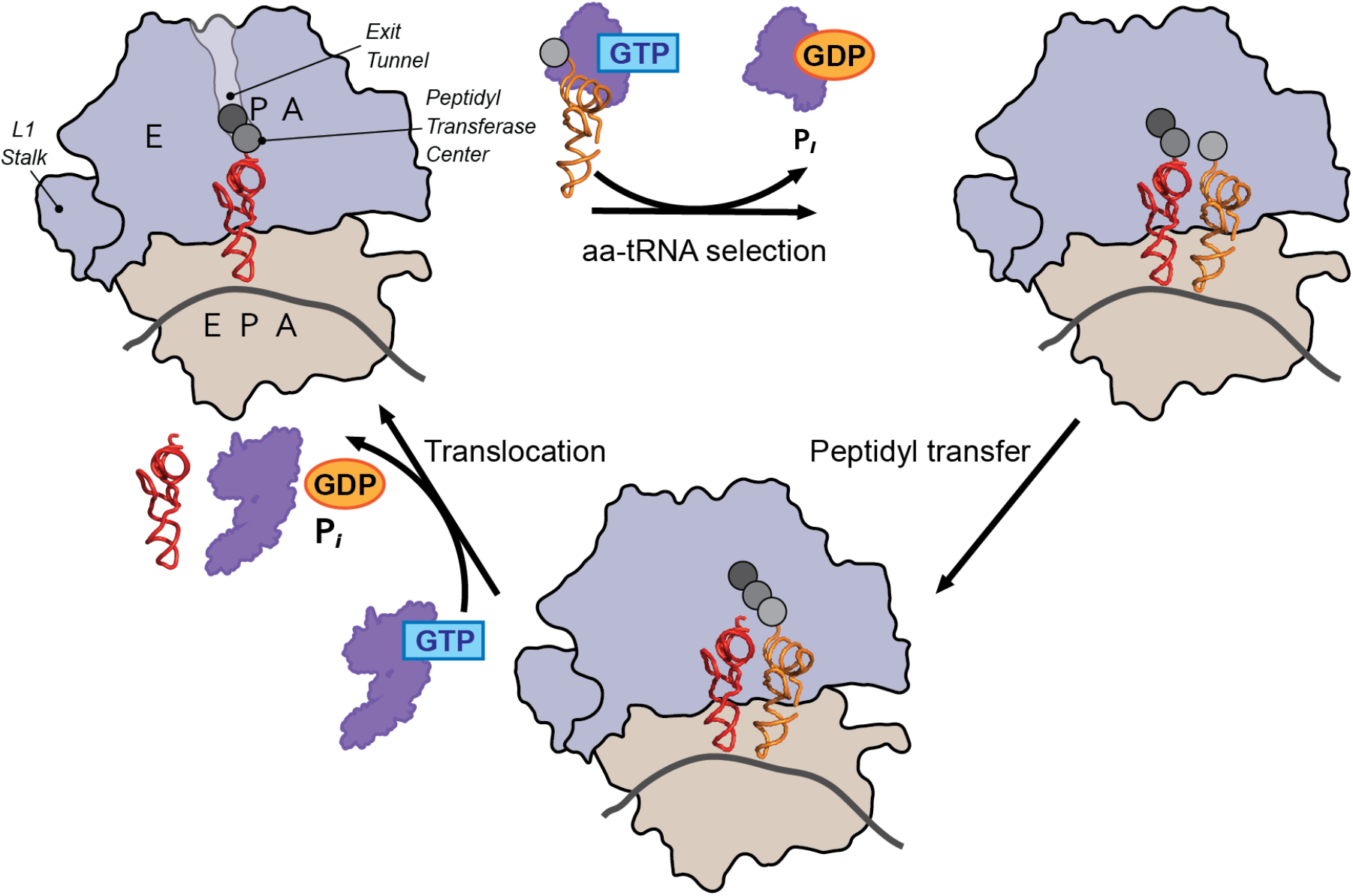
Overview of the translation elongation cycle. The elongation cycle consists of three major steps: aa-tRNA selection, peptidyl transfer, and translocation. See Section 3.2 for a detailed description of these steps.

### 3.3. Assaying the overall performance of ncAAs in polypeptide synthesis

Prior to assessing the details of any translation disorders that an ncAA might induce, a tripeptide synthesis reaction in which the ncAA is incorporated into the second residue position of the synthesized tripeptide is used to test the overall performance of an ncAA throughout all steps of the elongation cycle (Figure 2) (Englander et al., 2015; Fleisher et al., 2018). As detailed in the subsections that follow below, the tripeptide synthesis reaction is executed in four steps: (1) preparation of a 70S IC Mix, (2) preparation of a TC Mix, (3) preparation of an EF-G Mix, and (4) tripeptide synthesis *via* the addition of the TC Mix to a mixture of the 70S IC Mix and the EF-G Mix. Note that the protocols reported below for the preparation of the 70S IC Mix, TC Mix, and EF-G Mix and for performing tripeptide synthesis reactions are based on procedures that have been previously published (Blanchard, Kim, et al., 2004; Fei, 2010; Fei et al., 2010; Pavlov & Ehrenberg, 1996). Moreover, these protocols should serve as a general guide; for specific examples of the protocols we have used to prepare the 70S IC Mix, TC Mix, and EF-G Mix and perform tripeptide synthesis reactions in our mechanistic studies of ncAA incorporation, please refer to Effraim et al. (2009), Englander et al. (2015), Fleisher et al. (2018), and the smFRET experiments reported in Gamper et al. (2021).

**Figure 2:**
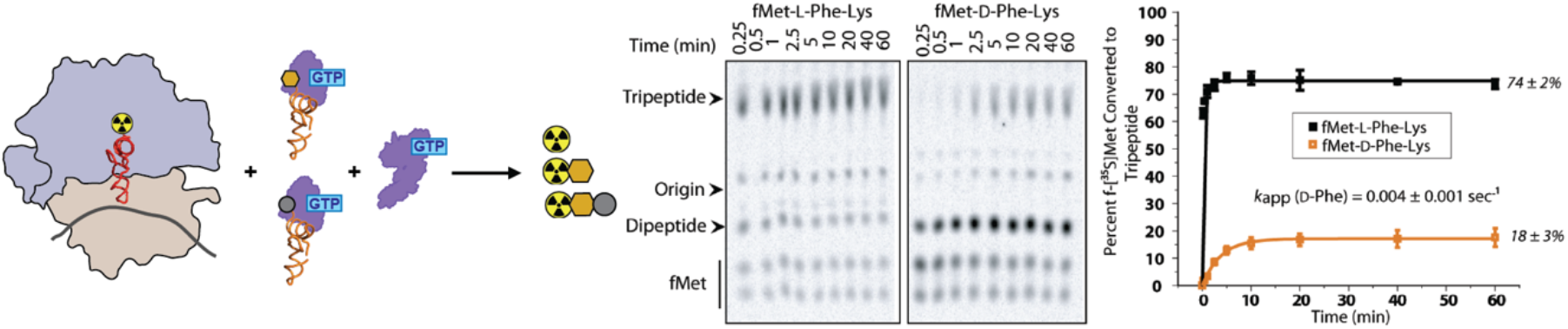
An assay that reports on the overall performance of ncAAs in polypeptide synthesis. The overall performance of an ncAA throughout all steps of the elongation cycle can be tested using a tripeptide synthesis assay in which the ncAA is incorporated into the second residue position. The ncAA is shown as a gold hexagon. See Section 3.3.6 for a detailed description of the assay. Data are shown for both an ncAA-tRNA (D-Phe-tRNA) and its corresponding cAA-tRNA (L-Phe-tRNA). The data and corresponding data figure panels are from Englander et al. (2015).

#### 3.3.1. Reagents, equipment, and general procedures

The following reagents and equipment were used for the tripeptide synthesis assays as well as for the other biochemical and smFRET assays described in Sections 3.4-3.8, unless otherwise specified. Chemical reagents, enzymes, and tRNAs were purchased from Sigma Aldrich unless otherwise specified. Depending on availability, it may be necessary to make appropriate substitutions for some of the equipment and instruments listed below.

##### Chemicals, solvents, and buffer salts

Tris-acetate (Tris-OAc)

Potassium chloride (KCl)

Ammonium acetate (NH_4_OAc)

Calcium acetate (Ca(OAc)_2_)

B-mercaptoethanol (BME)

Putrescine hydrochloride (putrescine-HCl)

Spermidine free base

Magnesium acetate (Mg(OAc)_2_)

Sodium hydroxide (NaOH)

Potassium hydroxide (KOH)

Guanosine 5’-triphosphate sodium salt hydrate (GTP) Ethylenediaminetetraacetic acid (EDTA)

Phosphoenol pyruvate (PEP)

Pyridine

Acetic acid

D-aa-containing peptides (chemically synthesized as described in Avins (2010) and Englander (2011))

Acetonitrile (MeCN)

##### Ribosomes, enzymes, mRNAs, and tRNAs

70S ribosomes (prepared as described in Section 3.1 and Fei et al. (2010))

Initiation Factor 1 (IF1) (prepared as described in Section 3.1 and Fei et al. (2010))

Initiation Factor 2 (IF2) (prepared as described in Section 3.1 and Fei et al. (2010))

Initiation Factor 3 (IF3) (prepared as described in Section 3.1 and Fei et al. (2010))

T4gp32_1–20_ mRNA (prepared as described in Section 3.1 and Fei et al. (2010))

f-[^35^S]Met-tRNA^fMet^ (prepared as described in Section 3.1 and Fei et al. (2010))

fMet-tRNA^fMet,^* (prepared as described in Section 3.1 and Fei et al. (2010))

Elongation Factor Tu (EF-Tu) (prepared as described in Section 3.1 and Fei et al. (2010))

Elongation Factor Ts (EF-Ts) (prepared as described in Section 3.1 and Fei et al. (2010))

Pyruvate kinase (PK)

2nd-position ncAA-tRNA (prepared as described in Section 2.3.4)

3^rd^-position cAA-tRNA (prepared as described in Section 3.1 and Fei et al. (2010))

Elongation Factor G (EF-G) (prepared as described in Section 3.1 and Fei et al. (2010))

##### Equipment and instruments

Eppendorf tubes

Ice bucket

VWR digital dry block heater (Model #949302)

Customized electrophoretic thin layer chromatography (eTLC) apparatus (assembly instructions kindly provided by Dr. Rachel Green, Johns Hopkins University)

VWR AccuPower power supply (Model #4000)

Phosphorimaging screen (GE Healthcare)

Phosphorimager (Typhoon FLA7000; GE Healthcare)

Reversed-phased Phenomenex Kinetex C18 column on a Waters 600 HPLC system or a reversed-phase Waters Xbridge C18 column on a Shimadzu LC-10AD^VP^ HPLC system*

##### Other supplies

Cellulose thin layer chromatography (TLC) plates (EMD)

Saran wrap

Image analysis software (ImageQuant, ImageJ, or similar)

Fiji image processing software*

*\*Needed for an alternative, HPLC-based, peptide product analysis method (Section 3.3.9) or preparation of ribosomal post-translocation (POST) complexes (Section 3.3.10)*

#### 3.3.2. Recipes

##### 5× Tris-Polymix Buffer at 25 mM Mg(OAc)_2_

250 mM Tris-OAc (pH_RT_ = 7.5)

500 mM KCl

25 mM NH_4_OAc

2.5 mM Ca(OAc)_2_

30 mM BME

25 mM putrescine-HCl

5 mM spermidine free base

25 mM Mg(OAc)_2_

Sterile filtered

If BME is omitted from the preparation, buffer lacking BME can be aliquoted and stored at –20°C, with BME being added just prior to use. Otherwise, if BME is included in the preparation, buffer containing BME should be made fresh just prior to use.

##### 5× Tris-Polymix Buffer at 0 mM Mg(OAc)_2_

250 mM Tris-OAc (pH_RT_ = 7.5)

500 mM KCl

25 mM NH_4_OAc

2.5 mM Ca(OAc)_2_

30 mM BME

25 mM putrescine-HCl

5 mM spermidine free base

Sterile filtered

If BME is omitted from the preparation, buffer lacking BME can be aliquoted and stored at –20°C, with BME being added just prior to use. Otherwise, if BME is included in the preparation, buffer containing BME should be made fresh just prior to use.

##### 50 mM GTP

50 mM GTP (pH_4°C_ = 7.0; titrated with 1 M NaOH)

Sterile filtered

Aliquoted and stored at –20 °C

##### TC Buffer

50 mM Tris-OAc (pH_RT_ = 7.5)

100 mM KCl

50 mM NH_4_OAc

0.5 mM Ca(OAc)_2_

0.1 mM EDTA

5 mM Mg(OAc)_2_

6 mM BME

Sterile filtered

If BME is omitted from the preparation, buffer lacking BME can be aliquoted and stored at –20°C, with BME being added just prior to use. Otherwise, if BME is included in the preparation, buffer containing BME should be made fresh just prior to use.

##### 300 mM PEP

300 mM PEP (pH_4°C_ = 7.0; titrated with 1 M KOH)

Sterile filtered

Aliquoted and stored at –20 °C

GTP Charging Mix

Employing TC Buffer as a diluent, use 50 mM GTP, 300 mM PEP, and PK to prepare a GTP Charging Mix composed of:

10 mM GTP

30 mM PEP

12.5 U/mL PK

Made fresh just prior to use

##### Pyridine Acetate Buffer

5% pyridine

20% acetic acid

pH_RT_ = 2.8

Stored at room temperature

#### 3.3.3. Preparation of the 70S IC Mix

a. When completed, the 70S IC Mix will be composed of 1.25 μM 70S ribosomes, 1.5 μM IF1, 1.5 μM IF2, 1.5 μM IF3, 4 μM T4gp32_1–20_ mRNA, 0.5 μM f-[^35^S]Met-tRNA^fMet^, and 1 mM GTP in Tris-Polymix Buffer at 5 mM Mg(OAc)_2_.
b. Depending on the targeted final volume of 70S IC Mix that will be necessary for the planned tripeptide synthesis reactions, calculate the number of moles of 70S ribosomes, IF1, IF2, IF3, GTP, f-[^35^S]Met-tRNA^fMet^, and T4gp32_1–20_ mRNA that will be needed for the 70S IC Mix.
c. Given their concentrations, calculate the volumes of the stock solutions of 70S ribosomes, IF1, IF2, IF3, 50 mM GTP, f-[^35^S]Met-tRNA^fMet^, and T4gp32_1–20_ mRNA that will be needed for the 70S IC Mix. Determine if it will be necessary to dilute any of the stock solutions in order to obtain volumes that can be accurately pipetted. If so, stock solutions can be diluted with Tris-Polymix Buffer at 5 mM Mg(OAc)_2_.
d. If prepared according to Fei et al. (2010) and Fei (2010), the stock solutions of 70S ribosomes and IF2 will contain 7.5 mM and 10 mM Mg(OAc)_2_, respectively, whereas none of the other stock solutions will contain any Mg^2+^. Given this as well as the volumes of stock solutions of 70S ribosomes and IF2 that will be added to the reaction mixture and the targeted final volume of 70S IC Mix, calculate the volumes of 5× Tris-Polymix Buffer at 25 mM Mg(OAc)_2_, 5× Tris-Polymix Buffer at 0 mM Mg(OAc)_2_, and nanopure water that will need to be added to the reaction mixture such that the targeted final volume of 70S IC Mix will be in Tris-Polymix Buffer at 5 mM Mg(OAc)_2_.
e. Given the calculations in Steps a-c, the appropriate volumes of 5× Tris-Polymix Buffer at 25 mM Mg(OAc)_2_, 5× Tris-Polymix Buffer at 0 mM Mg(OAc)_2_, nanopure water, 50 mM GTP, 70S ribosomes, IF1, IF2, and IF3 are added, in the listed order, to an Eppendorf tube that has been pre-cooled on ice.
f. The reaction mixture is carefully mixed by slowly pipetting up and down a few times and is incubated at 37 °C for 10 min. After the 10 min incubation, the reaction mixture is temporarily moved to room temperature to execute the next step.
g. Given the calculations in Steps a-c, the appropriate volume of T4gp32_1–20_ mRNA is then added to the reaction mixture.
h. The reaction mixture is again carefully mixed by slowly pipetting up and down a few times and is incubated at 37 °C for 10 min. After the 10 min incubation, the reaction mixture is temporarily moved to room temperature to execute the next step.
i. Given the calculations in Steps a-c, the appropriate volume of f-[^35^S]Met-tRNA^fMet^ is then added to the reaction mixture.
j. The reaction mixture is again carefully mixed by slowly pipetting up and down a few times and is incubated at 37 °C for 10 min. After this third 10 min incubation, the resulting 70S IC Mix is moved to ice, where it is stored until ready for use.
k. The 70S IC Mix is used without further purification and should be made fresh prior to each experiment.

#### 3.3.4. Preparation of the TC Mix

a. When completed, the TC Mix will be composed of 25 μM EF-Tu, 25 μM EF-Ts, 2.5 μM 2^nd^-position ncAA-tRNA, 2.5 μM 3^rd^-position cAA-tRNA, 1 mM GTP, 3 mM PEP, and 1.25 U/mL PK in TC Buffer.
b. Depending on the targeted final volume of TC Mix that will be necessary for the planned tripeptide synthesis reactions, calculate the number of moles of EF-Tu, EF-Ts, 2^nd^-position ncAA-tRNA, and 3^rd^-position cAA-tRNA that will be needed for the TC Mix.
c. Given their concentrations, calculate the volumes of the stock solutions of EF-Tu, EF-Ts, 2^nd^-position ncAA-tRNA, and 3^rd^-position cAA-tRNA that will be needed for the TC Mix. Determine if it will be necessary to dilute any of the stock solutions in order to obtain volumes that can be accurately pipetted. If so, stock solutions can be diluted with TC Buffer.
d. Given the volumes of stock solutions of EF-Tu, EF-Ts, 2^nd^-position ncAA-tRNA, and 3^rd^-position cAA-tRNA that will be added to the reaction mixture and considering that 1/10^th^ of the targeted final volume of TC Mix will come from the addition of GTP Charging Mix to the reaction mixture (see Step d), calculate the volume of TC Buffer that will need to be added to the reaction mixture to generate the targeted final volume of TC Mix.
e. Given the calculations in Steps a-c, the appropriate volumes of TC Buffer, GTP Charging Mix, EF-Tu, and EF-Ts are added, in the listed order, to an Eppendorf tube that has been pre-cooled on ice. Note that the volume of GTP Charging Mix added at this step will be 1/10^th^ of the targeted final volume of TC Mix (*i.e.*, such that the final concentrations of GTP, PEP, and PK in the TC Mix will be 1 mM GTP, 3 mM PEP, and 1.25 U/mL PK).
f. The reaction mixture is carefully mixed by slowly pipetting up and down a few times and is incubated at 37 °C for 1 min followed by an incubation on ice for 1 min. After the 1 min incubation on ice, the reaction mixture is temporarily kept on ice to execute the next step.
g. Given the calculations in Steps a-c, the appropriate volumes of the 2^nd^-position ncAA-tRNA and 3^rd^-position cAA-tRNA are then added to the reaction mixture.
h. The reaction mixture is again carefully mixed by slowly pipetting up and down a few times, is incubated at 37 °C for 1 min, and is then transferred to ice, where it is stored until ready for use.
i. The TC Mix is used without further purification and should be made fresh prior to each experiment.

#### 3.3.5. Preparation of the EF-G Mix

a. When completed, the EF-G Mix will be composed of 10 μM EF-G, 1 mM GTP, 3 mM PEP, and 1.25 U/mL PK in Tris-Polymix Buffer at 5 mM Mg(OAc)_2_.
b. Depending on the targeted final volume of EF-G Mix that will be necessary for the planned tripeptide synthesis reactions, calculate the number of moles of EF-G that will be needed for the EF-G Mix.
c. Given its concentration, calculate the volume of the stock solution of EF-G that will be needed for the EF-G Mix. Determine if it will be necessary to dilute the stock solution of EF-G in order to obtain a volume that can be accurately pipetted. If so, the stock solution can be diluted with Tris-Polymix Buffer at 5 mM Mg(OAc)_2_.
d. Given the volume of stock solution of EF-G that will be added to the reaction mixture and considering that 1/10^th^ of the targeted final volume of EF-G Mix will come from the addition of GTP Charging Mix to the reaction mixture (see Step d), calculate the volume of Tris-Polymix Buffer at 5 mM Mg(OAc)_2_ that will need to be added to the reaction mixture to generate the targeted final volume of EF-G Mix.
e. Given the calculations in Steps a-c, the appropriate volumes of Tris-Polymix Buffer at 5 mM Mg(OAc)_2_, GTP Charging Mix, and EF-G are added, in the listed order, to an Eppendorf tube that has been pre-cooled on ice. Note that the volume of GTP Charging Mix added at this step will be 1/10^th^ of the targeted final volume of TC Mix (*i.e.*, such that the final concentrations of GTP, PEP, and PK in the EF-G Mix will be 1 mM GTP, 3 mM PEP, and 1.25 U/mL PK).
f. The reaction mixture is carefully mixed by slowly pipetting up and down a few times, is incubated at 37 °C for 1 min, and is then transferred to ice, where it is stored until ready for use.
g. The EF-G Mix is used without further purification and should be made fresh prior to each experiment.

#### 3.3.6. Tripeptide synthesis reactions

a. When completed, the tripeptide synthesis reactions will be composed of 0.5 μM 70S ribosomes, 0.6 μM IF1, 0.6 μM IF2, 0.6 μM IF3, 1.6 μM T4gp32_1–20_ mRNA, 0.2 μM f-[^35^S]Met-tRNA^fMet^, 10 μM EF-Tu, 10 μM EF-Ts, 1 μM 2^nd^-position ncAA-tRNA, 1 μM 3^rd^-position cAA-tRNA, 2 μM EF-G, 1 mM GTP, 1.8 mM PEP, and 0.75 U/mL PK in Tris-Polymix Buffer at 5 mM Mg(OAc)_2_.
b. The Eppendorfs containing the 70 IC Mix (Section 3.3.3), TC Mix (Section 3.3.4), and EF-G Mix (Section 3.3.5) are incubated at 37 °C for 3 min prior to use.
c. The EF-G Mix is added to the 70S IC Mix in a 1:2 volume ratio (*e.g.*, 10 μL of the EF-G Mix is added to 20 μL of the 70S IC Mix) and the mixture is carefully mixed by slowly pipetting up and down a few times.
d. The tripeptide synthesis reaction is initiated by adding the TC Mix to the mixture of the EF-G Mix and the 70S IC Mix in a 2:3 ratio (*e.g.*, 20 μL of the TC Mix is added to the 30 μL of the mixture of the EF-G Mix and the 70S IC Mix).
e. The reaction is carefully mixed by slowly pipetting up and down a few times and is incubated at 37 °C for the desired timepoint(s).
f. At each desired timepoint, an aliquot of the reaction is collected and is quenched with KOH so as to achieve final concentration of 160 mM KOH.
g. The peptide products formed during a tripeptide synthesis reaction are then analyzed using eTLC (Section 3.3.7).

#### 3.3.7. eTLC analysis of tripeptide synthesis reaction products

Once aliquots for all desired timepoints from the tripeptide synthesis reactions have been collected and quenched, a fraction of each aliquot is spotted onto the center of a cellulose TLC plate and the spots are allowed to air-dry. The TLC plate is coated with a thin layer of Pyridine Acetate Buffer that is spread over the plate using a serological pipette, being careful to let the buffer wet the reaction spots by capillary action. The reaction products are separated using eTLC, in which an electric current at a given voltage is applied to the TLC plate in Pyridine Acetate Buffer (Englander, 2011; Weinger et al., 2004). The instructions for assembling a customized eTLC apparatus for implementing eTLC analysis were kindly provided in a personal communication from Dr. Rachel Green at Johns Hopkins University. The distance a molecule travels along a TLC plate, or how it partitions into the stationary and mobile phases, when placed in an electric field is directly proportional to its net charge (Fitzsimmons et al., 2001). Thus, the time required for appropriate separation of peptide products depends on the composition of the peptides. eTLCs for peptides with a smaller net charge or lower polar character can generally be run for 20-30 min at 1200 V, while those with a larger net charge or higher polar character can generally be run for 1 hr at 800 V (Englander et al., 2015; Fleisher et al., 2018). The voltage and run-time for tripeptide synthesis reactions can be adjusted according to the specific characteristics of the peptides being analyzed. As described in Section 2.3.4, eTLC plates are then air-dried, wrapped in Saran wrap, and exposed to a phosphorimaging screen overnight; the screen is imaged using a phosphorimager; and the eTLC spot intensities are quantified using ImageQuant, Image J, *etc*. The percent fMet converted to tripeptide for each timepoint is calculated as (*I*_tri_) / (*I*_fMet_+*I*_di_+*I*_tri_) × 100, where *I*_tri_, *I*_fMet_, and *I*_di_ are the intensities of the spots corresponding to the tripeptide product, the unreacted f-[^35^S]Met, and the dipeptide product, respectively. Experiments are typically executed in duplicate or triplicate, and the mean yield for each timepoint is plotted as a function of time with error bars representing either the standard error of the mean in the case of duplicate measurements or the standard deviation in the case of triplicate measurements. The mean yields as a function of time are then fit to a single-exponential function of the form y = y_0_ + A(e^−x/τ^), where y_0_ is the time offset, A is the amplitude of the change, and τ is the time constant (Effraim, 2010; Englander et al., 2015; Fleisher et al., 2018).

For reactions that are difficult to quantify using the above method due to poor separation between di- and tripeptide products, the data quantification method described by Schindelin et al. (2012) can be used. This method requires running a control, dipeptide synthesis reaction in which the 2^nd^-position cAA-tRNA is omitted from the reaction. Fiji image processing software is then used to generate an intensity profile for each column of an eTLC containing separated reaction products from the tripeptide synthesis reactions as well as the control, dipeptide synthesis reaction. The profiles of all columns from the plate are then aligned according to the position of the origin spot and the position of the spot corresponding to the unreacted f-[^35^S]Met. The column corresponding to the control, dipeptide synthesis reaction is then used to fit a single Gaussian distribution function so that the location and peak width for the dipeptide product can be determined. With this in hand, the columns containing both di- and tripeptide products can be fit to two Gaussian distribution functions using the linear least squares method. One of these functions is then fixed to the location and width of the single Gaussian function obtained for the dipeptide control column, whereas the width and location is allowed to vary for the other distribution. The height is allowed to vary for both functions. The percent dipeptide converted to tripeptide can then be estimated using the equation [A_tri_ / (A_di_ + A_tri_)] × 100, where A_di_ and A_tri_ correspond to the areas under the dipeptide and tripeptide Gaussian functions, respectively.

#### 3.3.8. Tripeptide synthesis competition assay

How well or poorly an ncAA-tRNA competes with its cAA-tRNA counterpart in protein synthesis can allow sensitive identification and initial characterization of any translation disorders exhibited by the ncAA-tRNA (Effraim et al., 2009; Englander et al., 2015; Fleisher et al., 2018). Tripeptide synthesis competition assays are executed and analyzed using a protocol identical to that described in Section 3.3.6, with the exception that the ncAA-tRNA and its cAA-tRNA counterpart, both cognate to the second codon position, are both included in the TC Mix at equal, 2.5 μM final concentrations and, consequently, in the tripeptide synthesis reaction at equal, 1 μM final concentrations.

#### 3.3.9. Chromatographic analysis for ascertaining chemical differences in the peptide products

Given that eTLC will not always be sensitive to the chemical differences between peptides containing an ncAA and their cAA counterparts (*e.g.*, between peptides with chemical differences in their peptide backbones, but otherwise identical amino acid sequences), reversed-phase C18 column chromatography in an HPLC system can be used to confirm that the synthesized polypeptide contains the ncAA. In our studies of D-aa incorporation by the TM, such an approach was used to verify the chirality of the peptide products obtained from the tripeptide synthesis assays using a solvent gradient unique to the peptides being analyzed; the solvent gradient is established using chemically synthesized, D-aa-containing, authentic, marker peptides (Avins, 2010; Englander, 2011). For example, to distinguish between fMet-D-Phe-Lys and fMet-L-Phe-Lys, in-house chemically synthesized, di- and tripeptide makers (Avins, 2010) were used to establish the following gradient for efficient peak separation between possible peptide products: 10% MeCN to 12% MeCN for 12 min, isocratic 20% MeCN from 12-15 min, and 20% MeCN to 35% MeCN from 15-58 min (Englander, 2011). This gradient was then used to confirm the chirality of the products obtained from our fMet-D-Phe-Lys and fMet-L-Phe-Lys tripeptide synthesis assays. For all racemization analyses of our tripeptide synthesis assays, unreacted f-[^35^S]Met and ribosome-synthesized peptide products were co-injected onto a reversed-phase C18 column together with the relevant marker peptides for peak identification (Englander et al., 2015). In general, this technique can be extended for use in distinguishing between peptides containing ncAAs that alter the chemistry of the peptide backbone (*e.g.*, D-aas, β-aas, *etc.*) and those containing their cAA counterparts with otherwise identical sequences by using in-house synthesized or commercially purchased marker peptides and establishing a solvent gradient for efficient peak separation.

#### 3.3.10. Preparation of a POST Complex

It is relatively straightforward to adapt the tripeptide synthesis assay protocols described in preceding sections to generate a ribosomal post-translocation (POST) complex that can be used in some of the biochemical assays described in the coming sections below. A 70S IC Mix, TC Mix, and EF-G Mix are prepared as described in Sections 3.3.3-3.3.5. Note that for the tripeptide synthesis assays, the limiting reactant is f-[^35^S]Met-tRNA^fMet^ (*i.e.*, we want to maximize the amount of f-[^35^S]Met-tRNA^fMet^ that is incorporated into 70S ICs). Thus, depending on the nature of the biochemical assay to be executed, it may be necessary to use non-radioactively labeled fMet-tRNA^fMet^, increase the concentration of this fMet-tRNA^fMet^ (up to 1 μM in the final tripeptide synthesis reaction), and/or decrease the concentration of a different reactant so as to make it the limiting reactant. Note also that, again depending on the nature of the biochemical assay to be executed, the TC Mix may use different concentrations of the 2^nd^-position ncAA-tRNA and/or the 3^rd^-position cAA-tRNA and/or may not include the 3^rd^-position cAA-tRNA and the EF-G Mix may or may not be necessary. Regardless of the choice of limiting reactant and absence or presence of the 3^rd^-position cAA-tRNA and EF-G Mix, a POST complex is prepared by following the protocol in Section 3.3.6 and adding TC Mix to a mixture of EF-G Mix, when included, and 70S IC Mix. These reactions are allowed to proceed long enough such that incorporation of the 2^nd^-position ncAA-tRNA and, when included, translocation and incorporation of the 3^rd^-position cAA-tRNA into the resulting POST complex have been maximized. For each 2^nd^-position ncAA-tRNA to be investigated, how long this takes needs to be empirically determined by executing a dipeptide synthesis assay (Section 3.4.1.) or a tripeptide synthesis assay (Section 3.3.6) and, if maximal translocation efficiency needs to be verified, a primer extension inhibition assay (Section 3.6.1).

### 3.4. Assaying delivery of ncAA-tRNAs to the A site and their performance as acceptors in the peptidyl transfer reaction

#### 3.4.1. Dipeptide synthesis assays

If the overall performance of an ncAA in a tripeptide synthesis assay is impaired, disorders in the early steps of the elongation cycle can be mechanistically characterized using a dipeptide synthesis assay. Specifically, dipeptide synthesis assays can be used to identify and characterize defects in how an ncAA-tRNA undergoes aa-tRNA selection and acts as an acceptor in the peptidyl transfer reaction, as such defects manifest as decreases in the yield and/or rate of dipeptide synthesis for reactions using ncAA-tRNAs relative to those using their cAA-tRNA counterparts (Figure 3A) (Effraim et al., 2009; Englander et al., 2015). Dipeptide synthesis assays are executed and analyzed analogously to the tripeptide synthesis assays described in Section 3.3.6, except that the 3^rd^-position aa-tRNA is omitted from the TC Mix and the percent fMet converted to dipeptide is quantified as (*I*_di_) / (*I*_fMet_ + *I*_di_) x 100. Reversed-phase C18 column chromatography and gradient development can be implemented as described in Section 3.3.9 using marker dipeptides.

**Figure 3:**
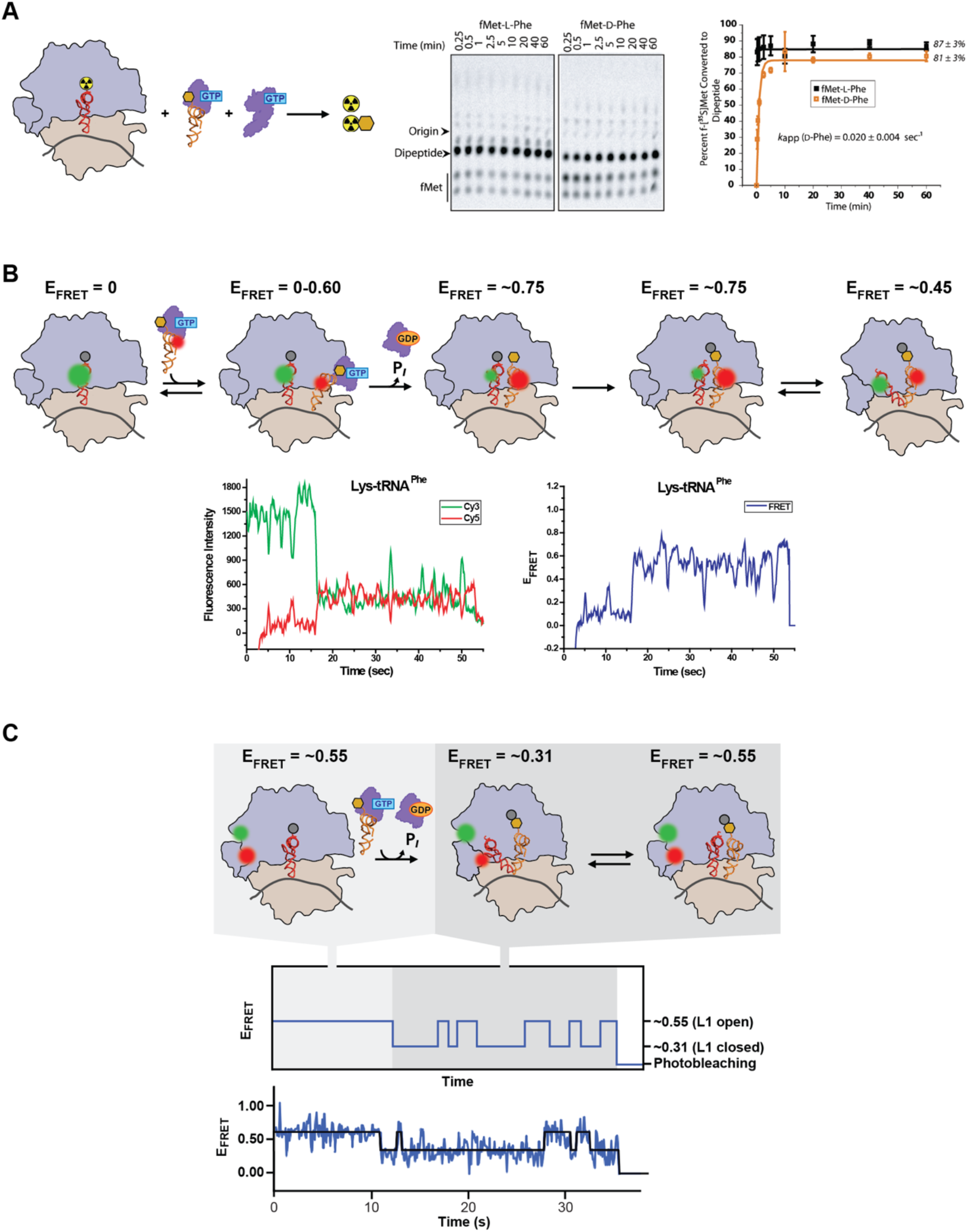
Assays that report on aa-tRNA selection and peptidyl transfer. The ability of an ncAA-tRNA to be delivered to the A site and serve as an acceptor in the peptidyl transfer reaction can be tested using: **(A)** a dipeptide synthesis assay, **(B)** a tRNA-tRNA smFRET assay, or **(C)** an L1-L9 smFRET assay. The ncAA is shown as a gold hexagon in all three panels. See Sections 3.4.1, 3.4.2, and 3.4.3 for detailed descriptions of the dipeptide synthesis-, tRNA-tRNA smFRET-, and L1-L9 smFRET assays, respectively. For the dipeptide synthesis assay, data are shown for both an ncAA-tRNA (D-Phe-tRNA) and its corresponding cAA-tRNA (L-Phe-tRNA). The data and corresponding data figure panels are from Englander et al. (2015). For the tRNA-tRNA assay, data are shown for a tRNA^Phe^ that has been misacylated with Lys. The data and corresponding data figure panels are from (Effraim et al., 2009). To clearly show the expected results for a cAA-tRNA that does not exhibit defects in aa-tRNA selection or peptidyl-transfer, the data shown for the L1-L9 smFRET assay correspond to a proline (Pro) that has been acylated onto a wildtype tRNA^Pro^ rather than onto a +1FS-inducing tRNA. The data and corresponding data figure panels are from Gamper et al. (2021).

#### 3.4.2. Dipeptide synthesis competition assay

Dipeptide synthesis competition assays report on how well or poorly an ncAA-tRNA competes with its cAA-tRNA counterpart in aa-tRNA selection and peptidyl transfer, as the yield of the ncAA-containing dipeptide will either be the same as or less than, respectively, that of the cAA-containing peptide (Effraim et al., 2009). These reactions are executed and analyzed analogously to those described in Section 3.4.1, except that the ncAA-tRNA and its cAA-tRNA counterpart, both cognate to the second codon position, are included at equal final concentrations of 1 μM each.

#### 3.4.3. aa-tRNA selection and peptidyl transfer smFRET assays

In addition to the ensemble biochemical assays described in Sections 3.4.1 and 3.4.2, we have also developed two smFRET assays that report on aa-tRNA selection and peptidyl transfer (Blanchard, Gonzalez, et al., 2004; Fei et al., 2009; Gonzalez Jr. et al., 2007; Kim et al., 2014), and have adapted these assays for use with ncAA-tRNAs (Figures 3B and C) (Effraim et al., 2009; Gamper et al., 2021). Biophysical smFRET experiments measure the efficiency of FRET (E_FRET_) between a FRET donor fluorophore and a FRET acceptor fluorophore that have been engineered into specific residue positions in a biomolecule or biomolecular complex of interest. The E_FRET_ depends, among other things, on the donor-acceptor distance, following the relation E_FRET_ = R_0_^6^ / R_0_^6^ + R^6^, where R_0_ is the donor-acceptor distance at which the E_FRET_ = 0.5 and is a characteristic constant for a particular donor-acceptor pair, and R is the donor-acceptor distance. In a typical biophysical smFRET experiment, E_FRET_ *versus* time trajectories (E_FRET_ trajectories) are measured for individual biomolecules or biomolecular complexes, thus allowing time-resolved changes in the donor-acceptor distance in individual biomolecules or biomolecular complexes to be directly observed. smFRET methods have been widely reviewed in the literature and, for appropriate background, the reader is referred to any of several comprehensive reviews (*e.g.*, (Joo et al., 2008; Lerner et al., 2018; Tinoco & Gonzalez, 2011).

For the smFRET studies described in this chapter, we use highly purified, fluorophore-labeled *E. coli* translation components prepared using the detailed protocols we have previously published in the *Methods in Enzymology* chapter by Fei et al. (2010). Briefly, tRNA^fMet^ that has been purchased or purified as described in Section 3.1 is labeled with a maleimide-derivatized, cyanine (Cy) 3 FRET donor fluorophore (Lumiprobe) at its naturally occurring 4-thiouridine at residue position 8. The resulting (Cy3)tRNA^fMet^ is methionylated and formylated as described in Section 3.1. Similarly, tRNA^Phe^ and tRNA^Lys^ that have been purchased or purified as described in Section 3.1 are labeled with *N*-hydroxysuccinimide ester (NHS)-derivatized, Cy5 FRET acceptor fluorophores (Lumiprobe) at their naturally occurring 3-(3-amino-3-carboxypropyl)uridine residues at position 47 and the resulting (Cy5)tRNA^Phe^ or (Cy5)tRNA^Lys^ are aminoacylated with the ncAA of interest as reported in Section 2.2. The 50S subunit is labeled with Cy3 and Cy5 by *in vitro* reconstituting recombinantly expressed, purified, and fluorophore-labeled variants of ribosomal proteins bL9 and uL1 into a 50S subunit lacking bL9 and uL1, having been purified from an *E. coli* strain lacking the genes encoding bL9 and uL1. Specifically, prior to *in vitro* reconstitution, bL9 is labeled with a maleimide-derivatized Cy3 at a single cysteine engineered into residue position 18 ((Cy3)bL9) and uL1 is labeled with a maleimide-derivatized Cy5 (Lumiprobe) at a single cysteine engineered into residue position 202 ((Cy5)uL1).

The first smFRET assay we have developed is a pre-steady state experiment in which the E_FRET_ trajectories report on aa-tRNA selection and peptidyl transfer using a FRET signal between fMet-(Cy3)tRNA^fMet^ in the P site of a 70S IC and a TC containing an aa-(Cy5)tRNA^Phe^ or aaAA-(Cy5)tRNA^Lys^ that is delivered to the A site (Figure 3B) (Blanchard, Gonzalez, et al., 2004; Effraim et al., 2009; Gonzalez Jr. et al., 2007). In this tRNA-tRNA smFRET experiment, the E_FRET_ value that is observed corresponds to the distance between our labeling positions on the P site-bound fMet-(Cy3)tRNA^fMet^ and the incoming aa-(Cy5)tRNA. A typical E_FRET_ trajectory thus begins at an E_FRET_ value of ∼0 before a TC interacts with the 70S IC and then transiently samples E_FRET_ values of ∼0 to ∼0.60 and returns to an E_FRET_ ∼0 several times as TCs non-productively and transiently sample the 70S IC. Ultimately, the E_FRET_ trajectory arrives at a longer-lived E_FRET_ value of ∼0.75 *via* a multi-step process as a TC productively delivers an aa-tRNA into the A site. Subsequently, the E_FRET_ trajectory exhibits a transition to an E_FRET_ value of ∼0.45, demonstrating that the aa-tRNA has served as an acceptor in the peptidyl transfer reaction and the newly deacylated tRNA in the P site and newly formed peptidyl-tRNA in the A site have adopted an alternative configuration as the resulting ribosomal pre-translocation (PRE) complex undergoes a transition to an alternative, well-characterized global conformational state. In the absence of EF-G, the E_FRET_ trajectory undergoes fluctuations between E_FRET_ values of ∼0.75 and ∼0.45, corresponding to the two alternative configurations the tRNAs adopt as the PRE complex fluctuates between its two well-characterized global conformational states.

To execute these tRNA-tRNA smFRET assays, a 70S IC Mix is prepared as described in Section 3.3.3, with two exceptions. The first exception is that 2 μM fMet-(Cy3)tRNA^fMet^ is substituted for 0.5 μM f-[^35^S]Met-tRNA^fMet^ such that the 70S ribosomes are now the limiting reagent. The second is that either a chemically synthesized, 5’-biotinylated T4gp32_1–9_ mRNA or an *in vitro* transcribed, T4gp32_1–20_ mRNA that has been pre-hybridized to a 3’-biotinylated DNA oligonucleotide complementary to the first 18 nucleotides of the T4gp32_1–20_ mRNA (Effraim et al., 2009; Gamper et al., 2021) (collectively referred to as Bio-T4gp32 mRNA) is substituted for T4gp32_1–20_ mRNA. 70S ICs from the resulting 70S IC Mix are purified using sucrose density gradient ultracentrifugation as described in Fei (2010). Briefly, the 70S IC Mix is first diluted five-fold using Tris-Polymix Buffer at 25 mM Mg(OAc)_2_ that is at 4 °C such that the final Mg(OAc)_2_ concentration is 20 mM. Up to 100 μL of diluted 70S IC Mix is then loaded onto the top of a 14 × 89 mm thin-wall, polypropylene, ultracentrifuge tube (Beckman) containing 12.8 mL of a 10-40% sucrose density gradient that has been prepared in Tris-Polymix Buffer at 20 mM Mg(OAc)_2_ using a gradient maker (Biocomp) and that has been pre-equilibrated to 4 °C. Gradients are ultracentrifuged at 25,000 RPM in a SW41 ultracentrifuge rotor (Beckman) for 12 hr at 4 °C. The ultracentrifuged gradients are then analyzed and fractionated using a gradient analyzer outfitted with a UV detector and a fraction collector (Biocomp) and fractions corresponding to the 70S IC are pooled, aliquoted, flash frozen in liquid nitrogen, and stored at –80 °C until further use. To investigate the incorporation of an ncAA-tRNA using this assay, a TC Mix is prepared as described in Section 3.3.4, with the exception that ncAA-(Cy5)tRNA is substituted for ncAA-tRNA.

smFRET experiments are executed in microfluidic flowcells that are part of five-flowcell ‘chips’ assembled using 1 × 3 in, 1 mm thick quartz microscope slides (G. Finkenbeiner) and 30 × 24 mm, #1.5, 0.16–0.19 mm thick borosilicate cover slips (VWR) as described in Fei (2010). Briefly, a set of five 0.75 mm holes are drilled along one edge of the central 1 in of the 3-in side of each slide using a 0.75 mm, diamond-coated drill bit (Starlite Industries) and a corresponding set of five holes are drilled along the other edge of the central 1 in of the 3 in side of the slide such that five pairs of holes along the 1 in side of the slide are created. The surfaces of the drilled slides and the coverslips are then cleaned, aminosilanized with Vectabond reagent (Vector Laboratories), and derivatized with a mixture of amine-reactive variants of polyethylene glycol (PEG) and biotinylated PEG (Bio-PEG) (mPEG-SPA or mPEG-SVA and Biotin-PEG-NHS or Biotin-PEG-SVA, respectively, from Laysan Bio). For each slide, a 1 in × ∼1 mm strip of double-sided tape is applied above and below each of the five pairs of holes along the 1-in side of the slide to define five flowcells with a hole at each end of each flowcell, a coverslip is placed on top of the double-sided tape to create the five flowcells, and epoxy is used to seal the edges of the five flowcells. Each of the resulting five flowcells can hold ∼7 μL.

Just prior to use, a flowcell is readied for experiments as described in (Fei, 2010). Briefly, the flowcell is rinsed by pipetting 200 μL of TP50 Buffer (10mM Tris-OAc (pH_RT_ = 7.5, where the ‘RT’ superscript refers to ‘room temperature’) and 50 mM KCl) into the hole at one end of the flowcell. To passivate the surfaces of the flowcell against non-specific binding of proteins and nucleic acids, 20 μL of Blocking Solution (10 μM Bovine Serum Albumin (BSA; Invitrogen) and 10 μM Blocking DNA in TP50 Buffer) is then pipetted into the flowcell and the flowcell is incubated for 10 min at room temperature. Blocking DNA is composed of a DNA strand with the sequence 5’-CGT TTA CAC GTG GGG TCC CAA GCA CGC GGC TAC TAG ATC ACG GCT CAG CT-3’ (IDT, Inc.) and its Watson-Crick complementary strand (IDT, Inc.) and is prepared by mixing the two strands to a final concentration of 100 μM each in 10 mM Tris-OAc (pH_RT_ = 7.5), heating the mixture to 95 °C for 2 min in a heating block, slow cooling the mixture by placing the heating block on top of the lab bench, and, when the temperature reaches 70 °C, adding 2 M KCl to a final concentration of 50 mM. Streptavidin is then bound to the Bio-PEGs located on the slide- and coverslip surfaces of the flowcell by pipetting 20 μL of 1 μM streptavidin in Blocking Solution into the flowcell and incubating the flowcell for an additional 10 min at room temperature. 200 μL of TP50 Buffer followed by 200 μL of Tris-Polymix Buffer at 5 mM Mg(OAc)_2_ are then pipetted into the flowcell to remove any unbound BSA, Blocking DNA, and streptavidin. An aliquot of the fMet-(Cy3)tRNA^fMet^- and Bio-mRNA-containing 70S ICs that has been purified by sucrose density gradient ultracentrifugation is then diluted to a final concentration of 50-100 pM 70S IC using Tris-Polymix Buffer at 5 mM Mg(OAc)_2_ and the 70S ICs are tethered to the streptavidin-bound Bio-PEGs by pipetting 20 μL of the diluted 70S ICs into the flowcell and incubating the flowcell for 5 min at room temperature. 200 μL of Imaging Buffer (Tris-Polymix Buffer at 5 mM Mg(OAc)_2_ supplemented with an Oxygen Scavenging System and a Triplet-State Quencher Cocktail) is then pipetted into the flowcell to remove any untethered 70S ICs. The Oxygen Scavenging System is composed of final concentrations of 1% (v/v) β-D-glucose, 25 U mL^−1^ glucose oxidase (Sigma, Inc.) and 250 U mL^−1^ catalase (Sigma, Inc.) and the Triplet-State Quencher Cocktail is composed of final concentrations of 1 mM 1,3,5,7-cyclooctatetraene (Aldrich) and 1 mM p-nitrobenzyl alcohol (Fluka).

smFRET imaging is accomplished using a laboratory-built, prism-based, total internal reflection fluorescence (TIRF) microscope that we have previously described (Fei, 2010; Fei et al., 2008). This microscope uses a 532 nm, diode-pumped, solid-state laser (Laser Quantum) as an excitation source and an electron-multiplying charged coupled device (EMCCD) camera (Andor iXon Ultra 888, from Oxford Instruments) as a wide-field detector. The EMCCD camera enables imaging of hundreds of individual 70S ICs as movies with a time resolution of 25–100 msec frame^−1^. Additionally, this microscope is outfitted with a syringe pump that can be connected to the inlet hole of a flowcell so as to allow stopped-flow delivery of TC Mix or a mixture of TC Mix and EF-G Mix to surface-tethered 70S ICs during imaging.

To initiate the tRNA-tRNA smFRET experiment, the stopped-flow syringe is loaded with 60 μL of a TC Mix that has been diluted to a final concentration of 10–100 nM aa-(Cy5)tRNA using Imaging Buffer. Once a field-of-view has been selected and the microscope has been focused, imaging commences and, 2 sec into the movie, the stopped-flow device delivers 50 μL of the diluted TC Mix to the surface-tethered 70S ICs. The interaction of single TCs with up to hundreds of individually spatially resolved, surface-tethered 70S ICs are then recorded as a collection of Cy3- and Cy5 fluorescence intensity *versus* time trajectories that can then be converted to E_FRET_ trajectories using E_FRET_ = I_Cy5_ / (I_Cy3_ + I_Cy5_), where I_Cy3_ and I_Cy5_ are the fluorescence intensities of Cy3 and Cy5, respectively, to calculate E_FRET_ at each timepoint. E_FRET_ trajectories can then be analyzed using methods we have previously published (Bronson et al., 2009; van de Meent et al., 2014) to determine the rates with which: (*i*) TCs non-productively sample the 70S IC; (*ii*) a TC productively delivers an ncAA-tRNA into the A site; (*iii*) the ncAA-tRNA serves as an acceptor in the peptidyl transfer reaction, and the tRNAs adopt an alternative configuration as the PRE complex undergoes a transition to its alternative, well-characterized global conformational state; and (*iv*) the tRNAs transition between the two alternative configurations they adopt as the PRE complex fluctuates between its two well-characterized global conformational states.

The second smFRET assay we have developed is a pre-steady state experiment in which the E_FRET_ trajectories report on aa-tRNA selection and peptidyl transfer using a FRET signal between (Cy3)bL9 and (Cy5)uL1 on the 50S subunit (Figure 3C) (Fei et al., 2009; Gamper et al., 2021; Kim et al., 2014). In this L1-L9 smFRET experiment, the E_FRET_ value that is observed corresponds to the distance between our labeling positions on (Cy3)bL9 and (Cy5)uL1. Importantly, (Cy5)uL1 is located within the ‘L1 stalk’ structural element of the 50S subunit that adopts ‘open’ or ‘closed’ conformations relative to (Cy3)bL9, which is located within the ‘core’ of the 50S subunit. Because we (Fei et al., 2009; Gamper et al., 2021; Kim et al., 2014) and others (Cornish et al., 2009) have previously shown that the L1 stalk transitions between its open and closed conformations at defined steps of the elongation cycle, the changes in the distance between our labeling positions on (Cy3)bL9 and (Cy5)uL1 that are captured by the E_FRET_ trajectories in this assay can be used to measure the kinetics of aa-tRNA selection and peptidyl transfer. Specifically, a typical E_FRET_ trajectory begins at an E_FRET_ value of ∼0.55 as the L1 stalk predominantly adopts the open conformation before a TC interacts with the 70S IC. Subsequently, the E_FRET_ trajectory exhibits a transition to an E_FRET_ value of ∼0.31, demonstrating that a TC has productively delivered an aa-tRNA into the A site, the aa-tRNA has served as an acceptor in the peptidyl transfer reaction, and the L1 stalk has closed as the PRE complex undergoes a transition to its alternative, well-characterized global conformational state. In the absence of EF-G, the E_FRET_ trajectory undergoes fluctuations between E_FRET_ values of ∼0.55 and ∼0.31, corresponding to the open and closed conformations that the L1 stalk adopts as the PRE complex fluctuates between its two well-characterized global conformational states.

L1-L9 smFRET assays are executed and analyzed in a manner analogous to that described above for the tRNA-tRNA smFRET assays with the exception that the 70S IC is prepared using (Cy3)bL9- and (Cy5)uL1-labeled 50S subunits and unlabeled fMet-tRNA^fMet^ and the TC is prepared using an unlabeled ncAA-tRNA. The resulting E_FRET_ trajectories can be analyzed to determine the rates with which: (*i*) a TC productively delivers an ncAA-tRNA into the A site, the ncAA-tRNA serves as an acceptor in the peptidyl transfer reaction, and the L1 stalk closes as the resulting PRE complex undergoes a transition to its alternative, well-characterized global conformational state and (*ii*) the L1 stalk fluctuates between the open and closed conformation it adopts as the PRE complex fluctuates between its two well-characterized global conformational states.

### 3.5. Assaying translocation of ncAA-containing peptidyl-tRNAs from the A site to the P site

#### 3.5.1. Primer extension inhibition, or ‘toeprinting’, assays

Assuming that an ncAA-tRNA can be successfully delivered to the A site and can act as an acceptor in the peptidyl transfer reaction, primer extension inhibition, or ‘toeprinting’, assays can be used to identify and characterize defects in how a PRE complex carrying an ncAA-containing peptidyl-tRNA in the A site undergoes translocation (Figure 4A) (Englander et al., 2015). In a toeprinting assay, reverse transcriptase (RT) is used to extend a radiolabeled DNA primer that has been pre-annealed to the mRNA downstream of a ribosomal complex assembled on the mRNA. RT is strongly blocked upon encountering the ribosomal complex, producing a cDNA primer extension product (*i.e.*, a ‘toeprint’) whose length, measured using denaturing PAGE (D-PAGE), marks the position of the ribosomal complex on the mRNA with single-nucleotide resolution. Toeprinting reactions executed on ribosomal complexes captured as they engage in peptide synthesis can therefore report on the extent and kinetics of translocation.

**Figure 4:**
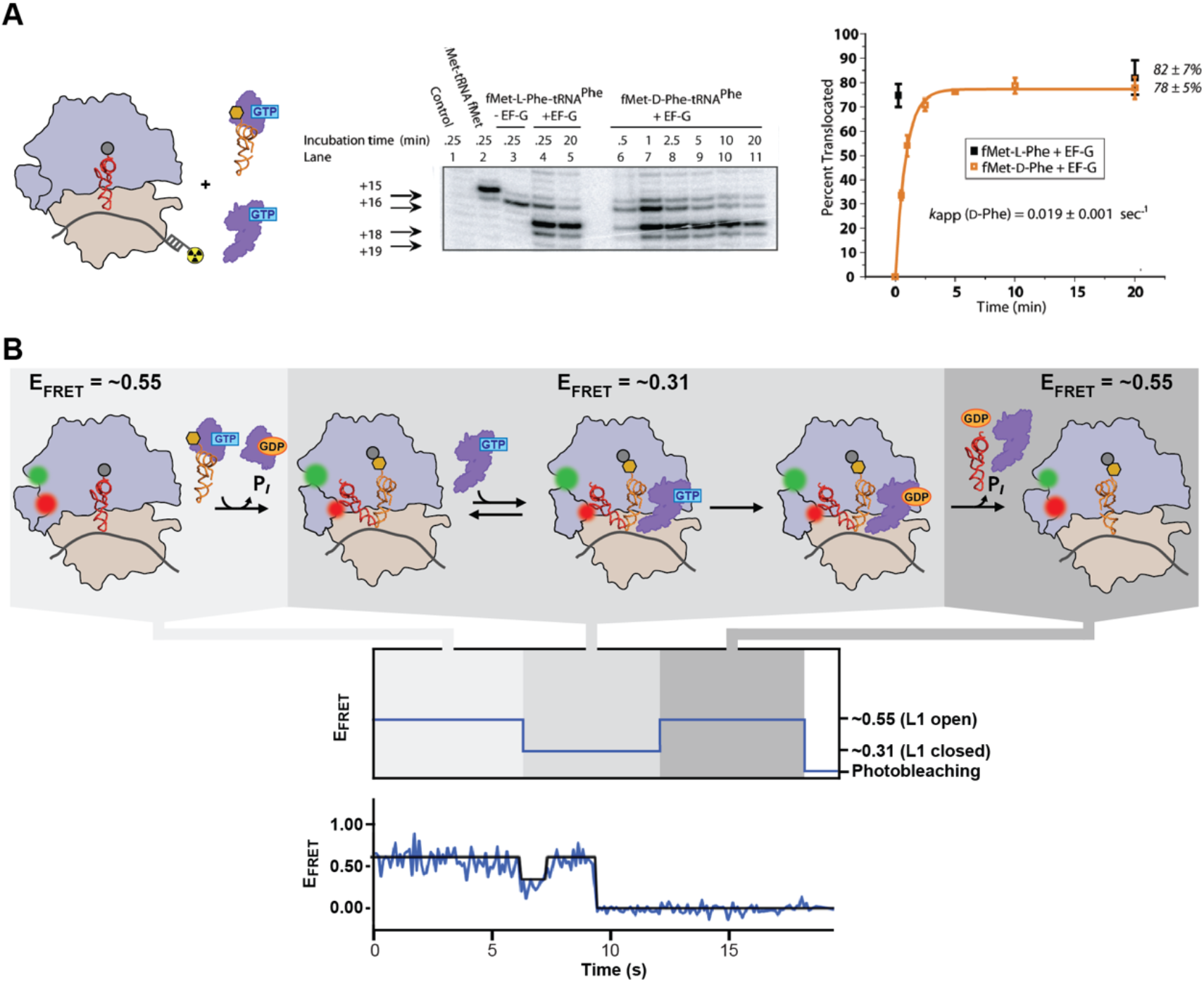
Assays that report on translocation. The ability of an ncAA-containing peptidyl-tRNA to be translocated from the A site to the P site can be tested using: **(A)** a primer extension inhibition, or ‘toeprinting’ assay or **(B)** an L1-L9 smFRET assay. The ncAA is shown as a gold hexagon in both panels. See Sections 3.5.1, and 3.5.2 for detailed descriptions of the toeprinting and L1-L9 smFRET assays, respectively. For the toeprinting assay, data are shown for both an ncAA-tRNA (D-Phe-tRNA) and its corresponding cAA-tRNA (L-Phe-tRNA). The data and corresponding data figure panels are from Englander et al. (2015). To clearly show the expected results for an cAA-containing peptidyl-tRNA that does not exhibit defects in translocation, the data shown for the L1-L9 smFRET assay correspond to a proline (Pro) that has been acylated onto a wildtype tRNA^Pro^ rather than onto a +1FS-inducing tRNA. The data and corresponding data figure panels are from Gamper et al. (2021).

It is important to use an mRNA that has as little intrinsic, thermodynamically stable secondary structure as possible along the segment that will be reverse transcribed; this minimizes the possibility of strong, mRNA secondary structure-induced RT stops that can make it difficult to interpret the results of the experiment. To this end, we have found that T4gp32_1–224_ mRNA contains less problematic secondary structure than T4gp32_1–20_ mRNA (Englander et al., 2015) and therefore use T4gp32_1–224_ mRNA in our toeprinting assays.

Radiolabeling the DNA primer (5′-TATTGCCATTCAGTTTAG-3′; (IDT, Inc.)) with γ[^32^P]ATP is carried out by combining 2.4 μM of the primer with 1.4 μM γ[^32^P]ATP (Perkin Elmer) and 0.5 U/μL T4 polynucleotide kinase (New England Biolabs) in Polynucleotide Kinase Buffer (New England Biolabs) and incubating for 30 min at 37 °C. The kinase is then inactivated by heating the reaction to 75 °C for 10 min and unincorporated radiolabeled nucleotides are subsequently removed by gel filtration through a G25 Sephadex spin column (GE Healthcare Life Sciences). The [^32^P]-labeled primer is annealed to the mRNA by combining 0.25 μM of the primer with 5 μM of the mRNA in 25 mM Tris-OAc (pH_RT_ = 7.0), incubating for 1.5 min at 90 °C, and slow cooling on the benchtop until the mixture cools to room temperature.

We typically use dipeptide synthesis reactions for toeprinting assays and carry these out as described in Section 3.4.1 with several exceptions. First, preparation of the 70S IC Mix is adjusted so that 1.25 μM fMet-tRNA^fMet^ is substituted for 0.5 μM f-[^35^S]Met-tRNA^fMet^ and 0.625 μM primer-annealed T4gp32_1–224_ mRNA is substituted for 4 μM T4gp32_1–20_ mRNA such that the primer is now the limiting reagent. Second, at each desired time point, an aliquot of the dipeptide synthesis reaction is quenched with 4× reaction volume of Toeprinting Mix (1.25 mM viomycin (BOC Sciences); 625 μM each dGTP, dCTP, and dTTP; and 2.2 mM dATP in 1.25× Tris-Polymix Buffer at 12.5 mM Mg(OAc)_2_) and transferred to ice. The antibiotic viomycin is included because it strongly inhibits EF-G-catalyzed translocation (Fredrick & Noller, 2003).

For each ncAA-containing peptidyl-tRNA and its corresponding cAA-containing peptidyl-tRNA to be tested, four dipeptide synthesis reactions are run. All four reactions are executed as described in the previous paragraph with the following exceptions. For the first reaction, Tris-Polymix Buffer at 5 mM Mg(OAc)_2_ is substituted for 70S ribosomes when preparing the 70S Mix. For the second reaction, Tris-Polymix Buffer at 5 mM Mg(OAc)_2_ is substituted for TC Mix and EF-G Mix. For the third reaction, Tris-Polymix Buffer at 5 mM Mg(OAc)_2_ is substituted for EF-G Mix. The fourth reaction is run as described in the previous paragraph, with no exceptions. Thus, the first, second, third, and fourth reactions will mark the positions of mRNA secondary structure-induced stops, the 70S IC, the PRE complex, and the POST complex, respectively.

mRNAs are then reverse-transcribed by adding avian myeloblastosis virus (AMV) RT (Promega) to a final concentration of 0.6 U/μL and incubating for 15 min at 37 °C. To determine where in the mRNA sequence the secondary structure-induced stops, 70S IC, PRE complex, and POST complex are located, we additionally execute Sanger sequencing of the mRNA. For the Sanger sequencing reactions, we use the same primer-annealed T4gp32_1–224_ mRNA and implement the reverse transcription exactly as for the dipeptide synthesis reactions, except that we adjust the Toeprinting Mix to omit viomycin and, for each Sanger sequencing reaction, include 25 μM of the dideoxynucleotide corresponding to the nucleotide to be sequenced.

The resulting 5’-[^32^P]-labeled cDNA products are then phenol extracted twice, chloroform extracted twice, and ethanol precipitated. Precipitated cDNA pellets are resuspended in Gel Loading Buffer (23 M formamide, 0.09% bromophenol blue, and 0.09% xylene cyanol) and the cDNA products are separated using 9% polyacrylamide D-PAGE. Similar to what is described in Section 2.3.4, the gel is then dried on a gel drier, wrapped in Saran wrap, and exposed to a phosphorimaging screen overnight; the screen is imaged using a phosphorimager; and the band intensities are quantified using ImageQuant, ImageJ, *etc*. Because the bands corresponding to the 70S IC, PRE complex, and POST complex are expected at the +15, +16, and +18/+19 mRNA nucleotide positions (where the positive numbers refer to the number of nucleotides downstream from the adenosine nucleotide of the AUG start codon), respectively, percent translocation at each time point is calculated as [(*I*_+18_ + *I*_+19_)/(*I*_+15_ + *I*_+16_ + *I*_+18_ + *I*_+19_)] × 100, where *I_+n_* is the intensity of the band corresponding to the +n nucleotide position. Occasionally, corrections to this calculation need to be implemented to normalize for intensity differences across the various lanes and/or to correct for misincorporation and translocation of additional ncAA-tRNAs. For examples of these types of corrections, please see Englander et al. (2015). As described in Section 3.3.7, experiments are executed in duplicate or triplicate; the mean yield for each timepoint is plotted as a function of time with error bars representing either the standard error or the standard deviation; and the mean yields as a function of time are fit to a single-exponential function of the form y = y_0_ + A(e^−x/τ^).

#### 3.5.2. Translocation smFRET assay

In addition to the toeprinting assay, the L1-L9 smFRET assay that we use for mechanistic studies of ncAA-tRNA selection and peptidyl transfer (see Section 3.4.3) can be easily extended to allow mechanistic studies of translocation (Fei et al., 2009; Kim et al., 2014). Recently, we have used this smFRET assay to study translocation of a PRE complex carrying a +1FS-inducing peptidyl-tRNA in the A site that we have shown can be used to incorporate an ncAA in response to a quadruplet codon (Figure 4B) (Gamper et al., 2021). These experiments are executed and analyzed in a manner analogous to those described in Section 3.4.3 for L1-L9 smFRET studies of ncAA-tRNA selection and peptidyl transfer with the exception that the 60 μL of 10–100 nM TC Mix that is loaded into the stopped-flow syringe is supplemented with EF-G Mix to a final, saturating concentration of 2 µM EF-G.

As was the case for the L1-L9 smFRET assay for aa-tRNA selection and peptidyl transfer, a typical E_FRET_ trajectory begins at an E_FRET_ value of ∼0.55 as the L1 stalk adopts the open conformation. Then, upon incorporation of the aa-tRNA into the A site, peptidyl transfer, and the subsequent rearrangement of the resulting PRE complex, the E_FRET_ trajectory undergoes a transition to an E_FRET_ value of ∼0.31 as the L1 stalk undergoes an open→closed transition. In the presence of saturating amounts of EF-G, however, the E_FRET_ trajectory remains at an E_FRET_ value of ∼0.31 for a short period of time as EF-G binds to the PRE complex and transiently stabilizes it in the global conformational state in which the L1 stalk is closed. The E_FRET_ trajectory then undergoes a transition back to an E_FRET_ value of ∼0.55 corresponding to opening of the L1 stalk as the PRE complex undergoes translocation and is converted to a POST complex. The resulting E_FRET_ trajectories can therefore be analyzed to determine the rates with which: (*i*) a TC productively delivers an ncAA-tRNA into the A site, the ncAA-tRNA serves as an acceptor in the peptidyl transfer reaction, and the L1 stalk closes as the resulting PRE complex undergoes a transition to its alternative, well-characterized global conformational state and (*ii*) the L1 stalk opens as the PRE complex undergoes translocation and is converted to a POST complex.

### 3.6. Assaying the stability of ncAA-containing peptidyl-tRNAs at the P site

Even if an ncAA-containing peptidyl-tRNA can be successfully translocated into the P site, it is possible for the peptidyl-tRNA to prematurely dissociate from the P site, particularly if the nascent polypeptide chain is short (Forster et al., 2003; Karimi et al., 1998). Such premature dissociation of an ncAA-containing peptidyl-tRNA from the P site can be identified and characterized using a nitrocellulose filter binding assay (Figure 5A) (Englander et al., 2015). In this assay, ribosomes and ribosome-bound tRNAs are retained on the negatively-charged nitrocellulose filters due to positively charged patches found on the solvent-exposed surfaces of the ribosomal proteins. In contrast, unbound tRNAs, which are negatively charged, will pass through the filter. Thus, quantitation of how much f-[^35^S]Met is retained on a filter *versus* how much passes through the filter as a function of time after using an f-[^35^S]Met-tRNA^fMet^-containing 70S IC to initiate a peptide synthesis reaction allows the extent and kinetics of premature dissociation of the ncAA-containing peptidyl-tRNA from the P site to be determined.

**Figure 5:**
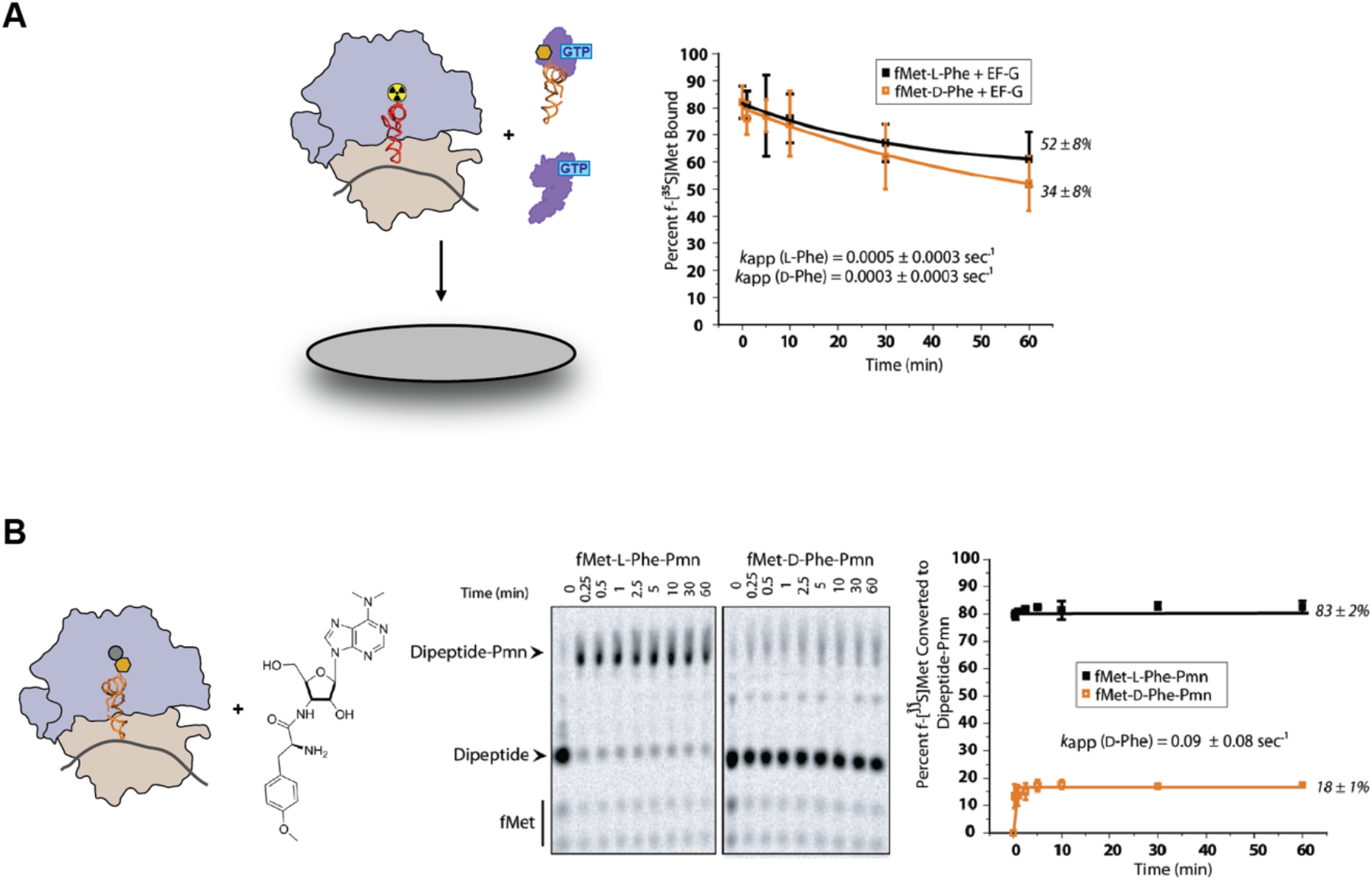
Assays that report on the stability of ncAA-containing peptidyl-tRNAs at the P site and their performance as donors in the peptidyl transfer reaction. **(A)** Following its translocation into the P site, the stability of an ncAA-containing peptidyl-tRNA at the P site can be tested using a nitrocellulose filter binding assay. The ncAA is shown as a gold hexagon. See Section 3.6 for a detailed description of this assay. Data are shown for both an ncAA-containing peptidyl-tRNA (fMet-D-Phe-tRNA) and its corresponding cAA-containing peptidyl-tRNA (fMet-L-Phe-tRNA). The data and corresponding data figure panels are from Englander et al. (2015). **(B)** The ability of an ncAA-containing peptidyl-tRNA that is stably bound at the P site to act as a donor in the peptidyl transfer reaction can be tested using a Pmn assay. The ncAA is shown as a gold hexagon. See Section 3.7 for a detailed description of this assay. Data are shown for both an ncAA-containing peptidyl-tRNA (fMet-D-Phe-tRNA) and its corresponding cAA-containing peptidyl-tRNA (fMet-L-Phe-tRNA). The data and corresponding data figure panels are from Englander et al. (2015).

Dipeptide synthesis reactions are executed as described in Section 3.4.1 with the exception that 0.5 μL of the mixture of EF-G Mix and 70S IC Mix is removed prior to the addition of TC Mix, is mixed with 49.5 μL of ice-cold Stop Buffer (50 mM Tris-HCl, pH_RT_ = 7.5, 1 M NH_4_Cl, 15 mM Mg(OAc)_2_), and is placed on ice. This will serve as the ‘zero’ time point of the reaction. Following addition of TC Mix to the mixture of EF-G Mix and 70S IC Mix, 0.5 μL of the dipeptide synthesis reaction is removed at each desired time point, mixed with 49.5 μL of ice-cold Stop Buffer, and placed on ice. For each Stop Buffer-treated time point, 30 μL of the reaction is applied to a nitrocellulose filter (Whatman) that has been pre-wetted with ice-cold Stop Buffer and placed over the well of a multi-well vacuum manifold (Millipore). An additional 10 μL of the reaction is applied to a second nitrocellulose filter that is not placed in the vacuum manifold. Vacuum is then applied to the manifold such that a flow rate of approximately 5 mL/min is established and maintained over the filters as they are washed extensively with ice-cold Stop Buffer. For each time point, the amount of f-[^35^S]Met retained on the filter that was exposed to vacuum and the filter that was not exposed to vacuum were determined by liquid scintillation counting (Perkin Elmer). For each time point, the percentage of f-[^35^S]Met retained on the nitrocellulose filter was calculated as (*C_+vac_*)/(3 × *C_–vac_*) × 100, where *C_+vac_* is the number of counts from the filter that was exposed to vacuum, *C_–vac_* is the number of counts from the filter that was not exposed to vacuum, and the factor of 3 is a normalization factor that accounts for the 3-fold larger volume of the reaction that was pipetted onto the filter that was exposed to vacuum relative to the one that was not exposed to vacuum. As described in Section 3.3.7, experiments are executed in duplicate or triplicate; the mean yield for each timepoint is plotted as a function of time with error bars representing either the standard error or the standard deviation; and the mean yields as a function of time are fit to a single-exponential function of the form y = y_0_ + A(e^−x/τ^).

### 3.7. Assaying the performance of ncAA-containing peptidyl-tRNAs as donors in the peptidyl transfer reaction

Assuming an ncAA-containing peptidyl-tRNA is successfully translocated into, and stably bound at, the P site, its ability to function as a donor in the peptidyl transfer reaction can be assessed using a puromycin (Pmn) assay (Figure 5B) (Englander et al., 2015). The antibiotic Pmn is an analog of the 3’ aminoacyl-adenosine end of an aa-tRNA that causes premature termination of polypeptide synthesis by binding to the A site, acting as an acceptor in the peptidyl transfer reaction, deacylating the peptidyl-tRNA at the P site, and ultimately dissociating from the A site, carrying the nascent polypeptide chain with it (Traut & Monro, 1964). As a minimal A-site substrate that does not undergo the rate-limiting, aa-tRNA selection steps that a full-length aa-tRNA would typically undergo, Pmn has been extensively used to probe the efficiency with which P site-bound peptidyl-tRNAs serve as donors in the peptidyl transfer reaction (Englander et al., 2015; Traut & Monro, 1964; Wohlgemuth et al., 2008).

We begin these assays by preparing a POST complex carrying an ncAA-containing peptidyl-tRNA at the P site as described in Section 3.3.10. To achieve this, we use a 70S IC Mix, TC Mix, and EF-G Mix prepared as described in Sections 3.3.3-3.3.5, with the exception that the 3^rd^-position cAA-tRNA is omitted from the TC Mix. As described in Section 3.3.10, POST complex formation reactions are incubated at 37 °C for as long as necessary to maximize incorporation of the 2^nd^-position ncAA-tRNA and subsequent translocation, as determined using a tripeptide synthesis assay (Section 3.3.6) and/or a primer extension assay (Section 3.6.1). As examples, to ensure 2^nd^-position D-aa-tRNAs are maximally incorporated and translocated during POST complex formation, we incubate the reactions for 10 min. In contrast, to ensure that their cAA-tRNA counterparts, L-aa-tRNAs, are maximally incorporated and translocated, we incubate them for only 2.5 min. Following this incubation, Pmn from a 100 mM stock solution prepared in Tris-Polymix Buffer is added to the POST complexes to a final Pmn concentration of 25 mM. Pmn reactions are quenched at the desired time points by adding KOH to a final concentration of 160 mM. As described in Section 3.3.7, reaction products are then separated by eTLC; eTLC plates are air-dried, wrapped in Saran wrap, and exposed to a phosphorimaging screen overnight; the screen is imaged using a phosphorimager; and the eTLC spot intensities are quantified using ImageQuant, Image J, *etc*. The percentage of f-[^35^S]Met converted to f-[^35^S]Met-X-Pmn, where X is the 2^nd^-position aa-tRNA, is quantified as (*I*_fM-X-Pmn_)/(*I*_fM_ + *I*_fM-X_ + *I*_fM-X-Pmn_) × 100, where *I*_fM-X-Pmn_, *I*_fM_, and *I*_fM-X_ are the eTLC spot intensities corresponding to the f-[^35^S]Met-X-Pmn and f-[^35^S]Met-X products and the unreacted f-[^35^S]Met, respectively. As described in Section 3.3.7, experiments are executed in duplicate or triplicate; the mean yield for each timepoint is plotted as a function of time with error bars representing either the standard error or the standard deviation; and the mean yields as a function of time are fit to a single-exponential function of the form y = y_0_ + A(e^−x/τ^).

### 3.8. Assaying ribosome structural changes imposed by ncAAs

Inhibition of translation by ncAAs can, at least in some cases, involve ncAA-mediated perturbation of the structure or structural dynamics of the ribosome. Nonetheless, several factors have until recently made structural studies of ncAA-containing ribosomal complexes challenging. As an example, samples are sometimes heterogeneous due to incomplete incorporation of the ncAA or to partitioning of the ncAA-containing ribosomal complexes into functional, and presumably, structural sub-populations (Englander et al., 2015; Fleisher et al., 2018). Such sample heterogeneity makes X-ray crystallography a poor choice. Although single-particle cryogenic electron microscopy (cryo-EM) should, in principle, enable classification of a structurally heterogeneous sample into a set of structural sub-populations, it is only recently that cryo-EM has attained the classification capabilities and high spatial resolution necessary to model and interpret the recorded Coulombic potential maps at near-atomic resolution. Such breakthroughs are now positioning single-particle cryo-EM to have a tremendous impact in the field (*vide infra*).

Because of the limitations described above and the fact that many of the perturbations of interest manifest as changes in the secondary structure or secondary structure dynamics of the ribosomal RNA (rRNA) component of the ribosome, chemical probing assays have served as a powerful approach for identifying and characterizing ncAA-mediated changes in the secondary structure or secondary structure dynamics of ncAA-containing ribosomal complexes. Even with the increasing feasibility of X-ray crystallographic and single-particle cryo-EM studies of ncAA-containing ribosomal complexes, chemical probing can serve as an excellent starting point for structural studies of ncAA-containing ribosomal complexes. In a rRNA chemical probing assay, the solvent accessibility of individual rRNA nucleotides is assessed by subjecting the rRNA to modification by different chemical probes that target either the bases or sugars of the nucleotides. By comparing the solvent accessibility of nucleotides in ncAA-containing ribosomal complexes with that of their cAA-containing counterparts, changes to rRNA secondary structure and/or secondary structure dynamics can be inferred. To this end, many RNA-reactive chemistries have been developed for chemical probing of RNA, including dimethyl sulfate (DMS), 1-cyclohexyl-3-(2-morpholinoethyl)carbodiimide metho-*p*-toluene sulfonate (CMCT), 3-ethoxy-a-ketobutyraldehyde (kethoxal), and selective 2’-hydroxyl acylation analyzed by primer extension (SHAPE) (Xu & Culver, 2009). In the course of our own work, we have used chemical probing with DMS to determine if and how an fMet-D-aa-tRNA bound at the P site of a POST complex alters the secondary structure or secondary structure dynamics of the PTC and nascent polypeptide exit tunnel entrance (ETE) sites within the 23S rRNA component of the 50S subunit (Figure 6). Based on this work, below we present a generalized protocol for chemical probing using DMS. We would like to note, however, that we strongly advocate the use of multiple probes for these types of studies and point the reader to many well-established, previously published protocols (Busan et al., 2019; Tijerina et al., 2007; Xu & Culver, 2009).

**Figure 6:**
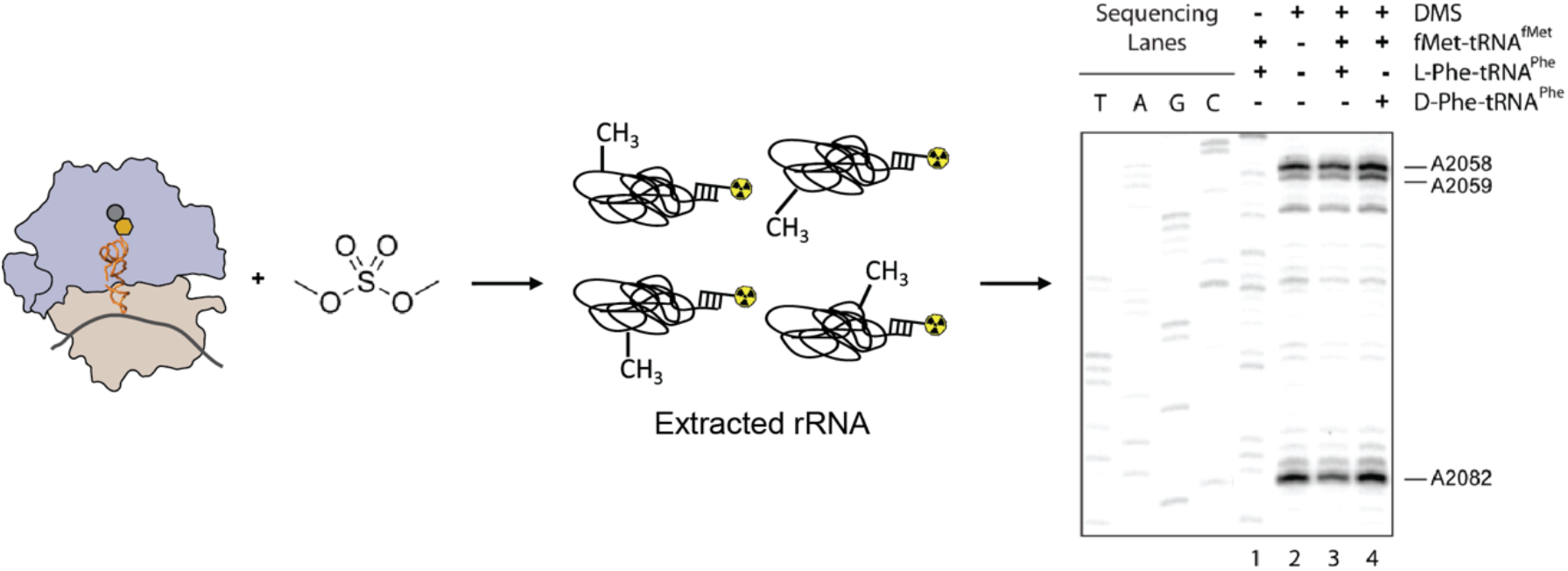
An assay that reports on ribosome structural changes imposed by ncAAs. The ability of an ncAA to inhibit translation by perturbing the structure or structural dynamics of the ribosome can be tested using a chemical probing assay. The ncAA is shown as a gold hexagon. See Section 3.8 for a detailed description of this assay. Data are shown for both an ncAA-containing peptidyl-tRNA (fMet-D-Phe-tRNA) and its corresponding cAA-containing peptidyl-tRNA (fMet-L-Phe-tRNA). The data and corresponding data figure panels are from Englander et al. (2015).

DMS is an alkylating agent that methylates the N1 position of adenine and, to a lesser extent, the N3 position of cytosine (Tijerina et al., 2007). Since these positions are involved in hydrogen bonding and/or have limited solvent accessibility when an adenine or cytosine is based-paired, only unpaired adenines and cytosines are robustly methylated by DMS. Methylation is achieved under conditions that result in ∼1 methylation event per rRNA molecule and the location of the methylated base in the rRNA is identified when it blocks the ability of an RT to transcribe past the methylated base. Specifically, a [^32^P]-labeled DNA primer is annealed to the rRNA extracted from a DMS-treated ribosomal complex and a cDNA is generated by primer extension using RT. When it encounters a methylated base, RT is blocked and can therefore no longer extend the cDNA, thereby generating a cDNA transcript of a specific length. cDNAs are then separated using D-PAGE alongside Sanger sequencing lanes to determine the exact position of the methylated base. Because a different base can be methylated in each rRNA molecule, the experiment generates a set of cDNAs and a corresponding set of bands on the gel that collectively map the location of all of the methylated bases. In addition, the more solvent accessible a particular base is, the more frequently it will be methylated and thus the more frequently its corresponding cDNA will appear in the set of cDNAs and the more intense its corresponding band will be on the gel. Comparing the location and intensities of the bands arising from different ribosomal complexes therefore allows changes in secondary structure or secondary structure dynamics to be detected with single-nucleotide resolution.

POST complexes carrying an ncAA-containing peptidyl-tRNA at the P site for these assays are prepared as described in Sections 3.3.10 and 3.7. This is accomplished using 70S IC Mix, TC Mix, and EF-G Mix prepared as described in Sections 3.3.3, 3.3.4, and 3.3.5, with the exceptions that 2 μM non-radiolabeled fMet-tRNA^fMet^ substitutes for the 1.25 μM f-[^35^S]Met-tRNA^fMet^ in the 70S IC Mix and the 3^rd^-position cAA-tRNA is omitted from the TC Mix. This ensures that the final concentration of fMet-tRNA^fMet^ in the POST complex formation reaction is 0.8 μM, with ribosomes being the limiting reagent. As described in Sections 3.3.10 and 3.7, POST complex formation reactions are incubated at 37 °C for as long as necessary to maximize incorporation of the 2^nd^-position ncAA-tRNA and subsequent translocation.

For each ncAA-containing peptidyl-tRNA to be tested, four POST complex formation reactions are run. All four reactions are executed as described in the previous paragraph with the following exceptions. For the first reaction, fMet-tRNA^fMet^ is excluded from the 70S IC Mix and ncAA-tRNA is excluded from the TC Mix so as to generate a vacant ribosomal complex containing no tRNAs. For the second and third reactions, a cAA-tRNA counterpart to the ncAA-tRNA to be tested is substituted for the ncAA-tRNA in the TC Mix. Note that the second reaction will not be treated with DMS (*vide infra*). The fourth reaction is run as described in the previous paragraph, with no exceptions. Thus, the first, second, third, and fourth reactions will map the locations of rRNA nucleotides that are accessible to DMS in the absence of tRNAs, result in secondary structure-induced RT stops, are protected from DMS in the presence of a cAA-containing peptidyl-tRNA, and are protected from DMS in the presence of an ncAA-containing peptidyl-tRNA, respectively.

Chemical probing of the POST complexes with DMS is then implemented using slight modifications of a previously published protocol (Stern et al., 1988). DMS is diluted 10-fold with DMS Buffer (80% ethanol in Tris-Polymix Buffer) immediately before use. 1 μL of the diluted DMS solution is added to the first, third, and fourth reactions described in the previous paragraph and 1 μL of DMS Buffer (*i.e.*, lacking DMS) is added to the second reaction described in the previous paragraph. All four samples are immediately incubated on ice for 45 min. Subsequently, 25 ng of glycogen (Ambion) are added as a carrier to all of the samples and the rRNA is precipitated with 95% ethanol. The precipitated rRNA pellets are then dissolved in 200 μL of Extraction Buffer (0.3 M sodium acetate (pH_RT_ = 5.2), 2.5 mM EDTA, and 0.5% sodium dodecyl sulfate (SDS)). The resuspended pellets are then extracted with phenol three times with vigorous agitation for 5 min followed by two subsequent extractions with chloroform with vigorous agitation for 3 min. This phenol-chloroform extraction procedure ensures that all ribosomal proteins are removed from the rRNA. Following extraction, the rRNA is ethanol precipitated a second time and the resulting pellets are dissolved in 15 μL of nanopure water.

A DNA primer complementary to the rRNA region of interest is [^32^P]-labeled as described in Section 3.5.1 (*e.g.*, to probe the PTC and ETE regions of 23S rRNA, we use a 17-nucleotide primer with the sequence 5′-CAAAGCCTCCCACCTAT-3′). The [^32^P]-labeled primer is annealed to the rRNA extracted from each POST complex by incubating a mixture of ∼1.0 pmol rRNA and 0.7 pmol of [^32^P]-labeled primer in a final volume of 10 μL of RT Buffer (25 mM Tris-HCl (pH_RT_ = 8.3), 40 mM KCl, and 5 mM MgCl_2_) for 5 min at 65 °C and then slow cooling to RT on the benchtop. Reverse transcription of the rRNA from each POST complex is then initiated by adding 10 μL of a solution that is 1.8 U/μL AMV RT and 1 mM in each dGTP, dCTP, dTTP, and dATP in RT Buffer to the 10 μL solution of [^32^P]-labeled primer-annealed rRNA in RT Buffer and the final, 20 μL reactions are incubated at 42 °C for 30 min.

To identify where in the rRNA sequence modified adenines and cytidines are located, we additionally execute Sanger sequencing of the rRNA. For the Sanger sequencing reactions, we use rRNA extracted from non-DMS-treated, vacant (*i.e.*, tRNA-free), 70S ribosomes in the same manner as we extracted the rRNA from the POST complexes. Annealing of the same [^32^P]-labeled primer to the rRNA extracted from the non-DMS-treated, vacant (*i.e.*, tRNA-free), 70S ribosomes and reverse transcription of the resulting [^32^P]-labeled primer-annealed rRNA are achieved identically to that of the rRNA extracted from the POST complexes, except that we adjust the solution containing the deoxynucleotides to include 25 μM of the dideoxynucleotide corresponding to the nucleotide to be sequenced.

The reverse transcription and Sanger sequencing reactions are quenched with 20 μL of Gel Loading Buffer and the cDNA products are separated using 7% polyacrylamide D-PAGE. As described in Section 3.5.1, the gel is dried, wrapped in Saran wrap, and exposed to a phosphorimaging screen overnight and the screen is imaged using a phosphorimager. We use the Semi-Automated Footprinting Analysis (SAFA) software program developed by Das et al. (2005) to quantify each gel using previously published protocols (Das et al., 2005; Englander et al., 2015). SAFA uses semi-automated routines to identify the lanes of a gel, identify the bands in each lane, assign each band in each lane to a nucleotide number, deconvolve any overlap from neighboring bands, correct for differences in loading amounts (using a normalization procedure developed by Takamoto et al. (2004) and implemented in SAFA), and quantify the intensity of each band. Experiments are typically executed in triplicate and from SAFA analysis of the three gels, we then determine the mean and standard deviation of the intensity of each band in the lanes corresponding to the POST complex carrying the ncAA-containing peptidyl-tRNA (*I*_ncAA_) and the cAA-containing peptidyl-tRNA (*I*_cAA_). For each nucleotide, *I*_ncAA_ and *I*_cAA_ values for the bands corresponding to that nucleotide are used to calculate *I*_cAA_/*I*_ncAA_. The entire set of *I*_cAA_/*I*_ncAA_ values are then used to calculate the average *I*_cAA_/*I*_ncAA_ (<*I*_cAA_/*I*_ncAA_>). Nucleotides exhibiting an *I*_cAA_/*I*_ncAA_ value that is some number of standard deviations above or below the <*I*_cAA_/*I*_ncAA_> value are then identified (*e.g.*, in our studies of D-aa incorporation, we used 2 standard deviations below or above the mean). These nucleotides are then interpreted as having significantly altered accessibilities to DMS, and therefore be involved in significantly altered secondary structures or secondary structure dynamics, in the POST complex carrying the ncAA-containing peptidyl-tRNA *versus* the POST complex carrying the cAA-containing peptidyl-tRNA.

## 4. Conclusions and Future Outlook

The toolbox of mechanistic assays described here has allowed us to explore the limits of misacylated cAA-tRNA (Effraim et al., 2009) and D-aa (Englander et al., 2015; Fleisher et al., 2018) incorporation by the TM as well as the limits of +1FS-mediated incorporation of ncAAs in response to quadruplet codons (Gamper et al., 2021). Notably, these assays are entirely general and can be used, adapted, or extended in order to identity and characterize the mechanistic underpinnings of inefficient incorporation of a broad range of ncAAs, thereby informing the ways in which the TM might be engineered to increase this efficiency.

Looking forward, we expect that emerging advances in the structural biology of engineered ribosomes (Schmied et al., 2018; Ward et al., 2019) and ribosomal complexes carrying ncAA-tRNAs (Melnikov et al., 2019) will be extended and, ultimately, expanded to include structural studies of other engineered ribosomes and TM components and of ribosomal complexes reflecting additional steps in the process of ncAA incorporation. Such studies promise to reveal the structural bases for the impaired or successful incorporation of ncAAs, information that could be invaluable for interpreting the results of mechanistic studies and informing efforts to engineer the TM. Likewise, we expect that increases in computational power will ultimately make it possible to run longer MD simulations on larger ribosomal complexes (Bock et al., 2018; Levi et al., 2019), developments that promise to bridge the gap between the all-atom, but static, structures generated by structural techniques such as X-ray crystallography and cryo-EM and the dynamic, but single-distance, structural rearrangements captured by smFRET. We expect that such MD simulations will continue to become important tools for helping to interpret biochemical, smFRET, and structural data on the incorporation of ncAAs; for providing roadmaps that enable engineering of the TM to improve the efficiency of ncAA incorporation; and for deciphering how engineered TM components facilitate the incorporation of ncAAs.

These are exciting times for the ncAA mutagenesis field. It is now becoming possible to combine high-resolution biochemical assays, smFRET experiments, structural studies, and MD simulations into a powerful framework that allows researchers to investigate the substrate specificity of the TM, identify and characterize the physical basis for limits in this specificity, and this information to engineer the TM, using either rational design or directed evolution approaches. We envision that such mechanistically informed engineering of the TM will unleash the full promise and potential of ncAA mutagenesis technology.

## Acknowledgements

We are indebted to Prof. Virginia W. Cornish, Prof. Thomas Leyh, and Prof. Ya-Ming Hou, all of whom have been essential collaborators and thought partners in our mechanistic studies of misacylated cAA-tRNA incorporation by the TM (V.W.C.), D-aa incorporation by the TM (V.W.C. and T.L.), and +1FS-mediated ncAA incorporation in response to quadruplet codons (Y.-M.H.). We thank Prof. Virginia W. Cornish, Prof. Thomas Leyh, Dr. Michael T. Englander, Dr. Philip R. Effraim, Dr. Joshua Avins, Dr. Gabriella Sanguinetti, Dr. Casey Brown, and Ms. Amanda Olivo-Ulichny for their contributions to our collaborative mechanistic studies of misacylated cAA-tRNA and/or D-aa incorporation by the TM and to the establishment of several of the biochemical assays described here; Prof. Rachel Green and Prof. Hani Zaher for kindly helping us establish the eTLC and nitrocellulose filter binding techniques in our laboratory; Prof. Joseph Puglisi, Prof. Steven Chu, Prof. Scott C. Blanchard, and Prof. Harold D. Kim for their contributions to our collaborative development of smFRET studies of the TM and the tRNA-tRNA smFRET signal; Dr. Jiangning Wang for adapting the tRNA-tRNA smFRET signal for smFRET studies of ncAA incorporation; Prof. Jingyi Fei and Dr. Haixing Li for their contributions to establishing and expanding the application of the L1-L9 smFRET signal for studies of ncAA incorporation; and Prof. Ya-Ming Hou, Prof. E. James Petersson, Prof. Gregor Blaha, Dr. Howard Gamper, Dr. Haixing Li, Dr. Isao Masuda, Dr. D. Miklos Robkis, Dr. Thomas Christian, and Dr. Adam Conn for their contributions to our collaborative mechanistic studies of +1FS-mediated ncAA incorporation in response to quadruplet codons. We fondly remember Prof. Klaus Schulten and we thank him and Dr. Bo Liu for the MD simulations they executed in support of our mechanistic studies of D-aa incorporation. This work was supported by National Institutes of Health (NIH) Grants GM090126 (to R.L.G.), GM084288 (to R.L.G.), GM119386 (to R.L.G), and GM137608 (to R.L.G.). This work was additionally funded by Burroughs Wellcome Fund Grant CABS 1004856 (to R.L.G.), National Science Foundation (NSF) Grant MCB 0644262 (to R.L.G.), as well as funding from Columbia University (to R.L.G.). R.C.F. was supported by the Training Program in Molecular Biophysics at Columbia University (T32 GM008281).

## References

Adamson, J. G., Hoang, T., Crivici, A., & Lajoie, G. A. (1992, Apr). Use of Marfey’s reagent to quantitate racemization upon anchoring of amino acids to solid supports for peptide synthesis. Anal Biochem, 202(1), 210–214. https://doi.org/10.1016/0003-2697(92)90229-z

Aleksashin, N. A., Leppik, M., Hockenberry, A. J., Klepacki, D., Vázquez-Laslop, N., Jewett, M. C., Remme, J., & Mankin, A. S. (2019, 2019/02/25). Assembly and functionality of the ribosome with tethered subunits. Nat Commun, 10(1), 930. https://doi.org/10.1038/s41467-019-08892-w

Aleksashin, N. A., Szal, T., d’Aquino, A. E., Jewett, M. C., Vázquez-Laslop, N., & Mankin, A. S. (2020, Apr 20). A fully orthogonal system for protein synthesis in bacterial cells. Nat Commun, 11(1), 1858. https://doi.org/10.1038/s41467-020-15756-1

Amiram, M., Haimovich, A. D., Fan, C., Wang, Y.-S., Aerni, H.-R., Ntai, I., Moonan, D. W., Ma, N. J., Rovner, A. J., Hong, S. H., Kelleher, N. L., Goodman, A. L., Jewett, M. C., Söll, D., Rinehart, J., & Isaacs, F. J. (2015, 2015/12/01). Evolution of translation machinery in recoded bacteria enables multi-site incorporation of nonstandard amino acids. Nature Biotechnology, 33(12), 1272–1279. https://doi.org/10.1038/nbt.3372

Avins, J. L. (2010). The Stereoselectivity of the Escherichia coli Protein Synthesis Machinery [PhD, Columbia University]. New York.

Bianco, A., Townsley, F. M., Greiss, S., Lang, K., & Chin, J. W. (2012, Sep). Expanding the genetic code of Drosophila melanogaster. Nat Chem Biol, 8(9), 748–750. https://doi.org/10.1038/nchembio.1043

Blanchard, S. C. (2002). Surface based translation: Single moelcule observation of ribosome activity Stanford University]. Satnford.

Blanchard, S. C., Gonzalez, R. L., Kim, H. D., Chu, S., & Puglisi, J. D. (2004, Oct). tRNA selection and kinetic proofreading in translation. Nat Struct Mol Biol, 11(10), 1008–1014.

Blanchard, S. C., Kim, H. D., Gonzalez, R. L., Puglisi, J. D., & Chu, S. (2004). tRNA dynamics on the ribosome during translation. Proceedings of the National Academy of Sciences of the United States of America, 101(35), 12893–12898. https://doi.org/10.1073/pnas.0403884101

Bock, L. V., Kolář, M. H., & Grubmüller, H. (2018, 2018/04/01/). Molecular simulations of the ribosome and associated translation factors. Current Opinion in Structural Biology, 49, 27–35. https://doi.org/10.1016/j.sbi.2017.11.003

Braun, T., Drescher, M., & Summerer, D. (2019). Expanding the Genetic Code for Site-Directed Spin-Labeling. International Journal of Molecular Sciences, 20(2). https://doi.org/10.3390/ijms20020373

Bronson, J. E., Fei, J., Hofman, J. M., Gonzalez Jr., R. L., & Wiggins, C. H. (2009). Learning rates and states from biophysical time series: A Bayesian approach to model selection and single-molecule FRET data. Biophysical Journal, 97, 3196–3205.

Busan, S., Weidmann, C. A., Sengupta, A., & Weeks, K. M. (2019, 2019/06/11). Guidelines for SHAPE Reagent Choice and Detection Strategy for RNA Structure Probing Studies. Biochemistry, 58(23), 2655–2664. https://doi.org/10.1021/acs.biochem.8b01218

Chen, S., Maini, R., Bai, X., Nangreave, R. C., Dedkova, L. M., & Hecht, S. M. (2017, 2017/10/11). Incorporation of Phosphorylated Tyrosine into Proteins: In Vitro Translation and Study of Phosphorylated IκB-α and Its Interaction with NF-κB. Journal of the American Chemical Society, 139(40), 14098–14108. https://doi.org/10.1021/jacs.7b05168

Chung, C. Z., Amikura, K., & Soll, D. (2020, Oct 19). Using Genetic Code Expansion for Protein Biochemical Studies. Frontiers in Bioengineering and Biotechnology, 8, Article 598577. https://doi.org/10.3389/fbioe.2020.598577

Cornish, P. V., Ermolenko, D. N., Staple, D. W., Hoang, L., Hickerson, R. P., Noller, H. F., & Ha, T. (2009, Feb 24). Following movement of the L1 stalk between three functional states in single ribosomes. Proceedings of the National Academy of Sciences of the United States of America, 106(8), 2571–2576.

Das, R., Laederach, A., Pearlman, S. M., Herschlag, D., & Altman, R. B. (2005, Mar). SAFA: semi-automated footprinting analysis software for high-throughput quantification of nucleic acid footprinting experiments. RNA, 11(3), 344–354. https://doi.org/11/3/344 [pii] 10.1261/rna.7214405

Datta, D., Wang, P., Carrico, I. S., Mayo, S. L., & Tirrell, D. A. (2002, 2002/05/01). A Designed Phenylalanyl-tRNA Synthetase Variant Allows Efficient in Vivo Incorporation of Aryl Ketone Functionality into Proteins. Journal of the American Chemical Society, 124(20), 5652–5653. https://doi.org/10.1021/ja0177096

Dedkova, L. M., Fahmi, N. E., Golovine, S. Y., & Hecht, S. M. (2003, 2003/06/01). Enhanced d-Amino Acid Incorporation into Protein by Modified Ribosomes. Journal of the American Chemical Society, 125(22), 6616–6617. https://doi.org/10.1021/ja035141q

Dedkova, L. M., Fahmi, N. E., Golovine, S. Y., & Hecht, S. M. (2006, 2006/12/01). Construction of Modified Ribosomes for Incorporation of d-Amino Acids into Proteins. Biochemistry, 45(51), 15541–15551. https://doi.org/10.1021/bi060986a

Dedkova, L. M., Fahmi, N. E., Paul, R., del Rosario, M., Zhang, L., Chen, S., Feder, G., & Hecht, S. M. (2012, 2012/01/10). β-Puromycin Selection of Modified Ribosomes for in Vitro Incorporation of β-Amino Acids. Biochemistry, 51(1), 401–415. https://doi.org/10.1021/bi2016124

Dedkova, L. M., & Hecht, S. M. (2019, 2019/04/24). Expanding the Scope of Protein Synthesis Using Modified Ribosomes. Journal of the American Chemical Society, 141(16), 6430–6447. https://doi.org/10.1021/jacs.9b02109

Desai, B. J., & Gonzalez, R. L. (2020, 2020/10/01). Multiplexed genomic encoding of non-canonical amino acids for labeling large complexes. Nature Chemical Biology, 16(10), 1129–1135. https://doi.org/10.1038/s41589-020-0599-5

Drienovská, I., & Roelfes, G. (2020, 2020/03/01). Expanding the enzyme universe with genetically encoded unnatural amino acids. Nature Catalysis, 3(3), 193–202. https://doi.org/10.1038/s41929-019-0410-8

Dunkelmann, D. L., Willis, J. C. W., Beattie, A. T., & Chin, J. W. (2020, 2020/06/01). Engineered triply orthogonal pyrrolysyl–tRNA synthetase/tRNA pairs enable the genetic encoding of three distinct non-canonical amino acids. Nature Chemistry, 12(6), 535–544. https://doi.org/10.1038/s41557-020-0472-x

Dupasquier, M., Kim, S., Halkidis, K., Gamper, H., & Hou, Y. M. (2008, Jun 6). tRNA integrity is a prerequisite for rapid CCA addition: implication for quality control. J Mol Biol, 379(3), 579–588. https://doi.org/10.1016/j.jmb.2008.04.005

Effraim, P. R. (2010). Revisiting the adaptor hypothesis: Studies of ribosomal selection of misacylated tRNAs (Publication Number 3420785) [Ph.D., Columbia University]. Dissertations & Theses @ Columbia University; ProQuest Dissertations & Theses Global. Ann Arbor.

Effraim, P. R., Wang, J., Englander, M. T., Avins, J. L., Leyh, T. S., Gonzalez, R. L., & Cornish, V. W. (2009). Natural amino acids do not require their native tRNAs for efficient selection by the ribosome. Nature Chemical Biology, 5(2009), 947–953.

Englander, M. T. (2011). The Ribosome Discriminates the Structure of the Amino Acid at its Peptidyl-Transferase Center [PhD, Columbia University]. New York.

Englander, M. T., Avins, J. L., Fleisher, R. C., Liu, B., Effraim, P. R., Wang, J., Schulten, K., Leyh, T. S., Gonzalez, R. L., & Cornish, V. W. (2015). The ribosome can discriminate the chirality of amino acids within its peptidyl-transferase center. Proceedings of the National Academy of Sciences, 112(19), 6038–6043. https://doi.org/10.1073/pnas.1424712112

Ernst, R. J., Krogager, T. P., Maywood, E. S., Zanchi, R., Beránek, V., Elliott, T. S., Barry, N. P., Hastings, M. H., & Chin, J. W. (2016, Oct). Genetic code expansion in the mouse brain. Nat Chem Biol, 12(10), 776–778. https://doi.org/10.1038/nchembio.2160

Fan, C., Xiong, H., Reynolds, N. M., & Söll, D. (2015). Rationally evolving tRNAPyl for efficient incorporation of noncanonical amino acids. Nucleic Acids Res, 43(22), e156–e156. https://doi.org/10.1093/nar/gkv800

Fei, J. (2010). Coupling of ribosome and tRNA dynamics during protein synthesis [PhD, Columbia University]. New York.

Fei, J., Bronson, J. E., Hofman, J. M., Srinivas, R. L., Wiggins, C. H., & Gonzalez, R. L., Jr. (2009, Sep 15). Allosteric collaboration between elongation factor G and the ribosomal L1 stalk directs tRNA movements during translation. Proc. Natl. Acad. Sci. USA, 106(37), 15702–15707. https://doi.org/0908077106 [pii] 10.1073/pnas.0908077106

Fei, J., Kosuri, P., MacDougall, D. D., & Gonzalez, R. L., Jr. (2008, May 9). Coupling of ribosomal L1 stalk and tRNA dynamics during translation elongation. Mol Cell, 30(3), 348–359.

Fei, J., Wang, J., Sternberg, S. H., MacDougall, D. D., Elvekrog, M. M., Pulukkunat, D. K., Englander, M. T., & Gonzalez, R. L. (2010). A Highly Purified, Fluorescently Labeled In Vitro Translation System for Single-Molecule Studies of Protein Synthesis. In N. G. Walter (Ed.), Methods in Enzymology (Vol. 472, pp. 221–259). Academic Press. https://doi.org/10.1016/S0076-6879(10)72008-5

Fitzsimmons, T. L., Fisher, T. L., & Reingold, I. D. (2001, 2001/09/01). Thin-Layer Electrophoresis. Journal of Chemical Education, 78(9), 1241. https://doi.org/10.1021/ed078p1241

Fleisher, R. C., Cornish, V. W., & Gonzalez, R. L. (2018, 2018/07/24). d-Amino Acid-Mediated Translation Arrest Is Modulated by the Identity of the Incoming Aminoacyl-tRNA. Biochemistry, 57(29), 4241–4246. https://doi.org/10.1021/acs.biochem.8b00595

Forster, A. C., Tan, Z., Nalam, M. N., Lin, H., Qu, H., Cornish, V. W., & Blacklow, S. C. (2003, May 27). Programming peptidomimetic syntheses by translating genetic codes designed de novo. Proc Natl Acad Sci U S A, 100(11), 6353–6357.

Fredrick, K., & Noller, H. F. (2003, May 16). Catalysis of ribosomal translocation by sparsomycin. Science, 300(5622), 1159–1162. http://www.ncbi.nlm.nih.gov/entrez/query.fcgi?cmd=Retrieve&db=PubMed&dopt=Citation&list_uids=12750524

Fried, S. D., Schmied, W. H., Uttamapinant, C., & Chin, J. W. (2015). Ribosome Subunit Stapling for Orthogonal Translation in E. coli. Angew Chem Int Ed Engl, 54(43), 12791–12794. https://doi.org/10.1002/anie.201506311

Gamper, H., Li, H., Masuda, I., Miklos Robkis, D., Christian, T., Conn, A. B., Blaha, G., Petersson, E. J., Gonzalez, R. L., & Hou, Y.-M. (2021, 2021/01/12). Insights into genome recoding from the mechanism of a classic +1-frameshifting tRNA. Nat Commun, 12(1), 328. https://doi.org/10.1038/s41467-020-20373-z

Gonzalez Jr., R. L., Chu, S., & Puglisi, J. D. (2007, Dec 2007). Thiostrepton inhibition of tRNA delivery to the ribosome. RNA, 13(12), 2091–2097.

Goodlett, D. R., Abuaf, P. A., Savage, P. A., Kowalski, K. A., Mukherjee, T. K., Tolan, J. W., Corkum, N., Goldstein, G., & Crowther, J. B. (1995, 1995/07/21/). Peptide chiral purity determination: hydrolysis in deuterated acid, derivatization with Marfey’s reagent and analysis using high-performance liquid chromatography-electrospray ionization-mass spectrometry. Journal of Chromatography A, 707(2), 233–244. https://doi.org/10.1016/0021-9673(95)00352-N

Hammerling, M. J., Krüger, A., & Jewett, M. C. (2020, Feb 20). Strategies for in vitro engineering of the translation machinery. Nucleic Acids Res, 48(3), 1068–1083. https://doi.org/10.1093/nar/gkz1011

Heckler, T. G., Roesser, J. R., Xu, C., Chang, P. I., & Hecht, S. M. (1988, Sep 20). Ribosomal binding and dipeptide formation by misacylated tRNA(Phe),S. Biochemistry, 27(19), 7254–7262. https://doi.org/10.1021/bi00419a012

Hou, Y. M., Gamper, H., & Yang, W. (2015, Apr). Post-transcriptional modifications to tRNA--a response to the genetic code degeneracy. RNA, 21(4), 642–644. https://doi.org/10.1261/rna.049825.115

Isaacs, F. J., Carr, P. A., Wang, H. H., Lajoie, M. J., Sterling, B., Kraal, L., Tolonen, A. C., Gianoulis, T. A., Goodman, D. B., Reppas, N. B., Emig, C. J., Bang, D., Hwang, S. J., Jewett, M. C., Jacobson, J. M., & Church, G. M. (2011). Precise Manipulation of Chromosomes in Vivo Enables Genome-Wide Codon Replacement. Science, 333(6040), 348. https://doi.org/10.1126/science.1205822

Jelenc, P. C., & Kurland, C. G. (1979). Nucleoside triphosphate regeneration decreases the frequency of translation errors. Proceedings of the National Academy of Sciences, 76(7), 3174. https://doi.org/10.1073/pnas.76.7.3174

Joo, C., Balci, H., Ishitsuka, Y., Buranachai, C., & Ha, T. (2008). Advances in single-molecule fluorescence methods for molecular biology. Annual Review of Biochemistry, 77, 51–76.

Karimi, R., Pavlov, M. Y., Heurgué-Hamard, V., Buckingham, R. H., & Ehrenberg, M. (1998, Aug 14). Initiation factors IF1 and IF2 synergistically remove peptidyl-tRNAs with short polypeptides from the P-site of translating Escherichia coli ribosomes. J Mol Biol, 281(2), 241–252. https://doi.org/10.1006/jmbi.1998.1953

Katoh, T., Iwane, Y., & Suga, H. (2017). Logical engineering of D-arm and T-stem of tRNA that enhances d-amino acid incorporation. Nucleic Acids Res, 45(22), 12601–12610. https://doi.org/10.1093/nar/gkx1129

Katoh, T., Sengoku, T., Hirata, K., Ogata, K., & Suga, H. (2020, 2020/11/01). Ribosomal synthesis and de novo discovery of bioactive foldamer peptides containing cyclic β-amino acids. Nat Chem, 12(11), 1081–1088. https://doi.org/10.1038/s41557-020-0525-1

Katoh, T., & Suga, H. (2018, 2018/09/26). Ribosomal Incorporation of Consecutive β-Amino Acids. Journal of the American Chemical Society, 140(38), 12159–12167. https://doi.org/10.1021/jacs.8b07247

Katoh, T., Tajima, K., & Suga, H. (2017, Jan 19). Consecutive Elongation of D-Amino Acids in Translation. Cell Chem Biol, 24(1), 46–54. https://doi.org/10.1016/j.chembiol.2016.11.012

Kazayama, A., Yamagami, R., Yokogawa, T., & Hori, H. (2015, May). Improved solid-phase DNA probe method for tRNA purification: large-scale preparation and alteration of DNA fixation. J Biochem, 157(5), 411–418. https://doi.org/10.1093/jb/mvu089

Kim, H. K., Liu, F., Fei, J., Bustamante, C., Gonzalez, R. L., Jr., & Tinoco, I., Jr. (2014, Apr 15). A frameshifting stimulatory stem loop destabilizes the hybrid state and impedes ribosomal translocation. Proc Natl Acad Sci U S A, 111(15), 5538–5543. https://doi.org/10.1073/pnas.1403457111

Kochhar, S., Mouratou, B., & Christen, P. (2000). Amino acid analysis by high-performance liquid chromatography after derivatization with 1-fluoro-2,4-dinitrophenyl-5-L-alanine amide (Marfey’s reagent). Methods Mol Biol, 159, 49–54. https://doi.org/10.1385/1-59259-047-0:049

Lajoie, M. J., Rovner, A. J., Goodman, D. B., Aerni, H.-R., Haimovich, A. D., Kuznetsov, G., Mercer, J. A., Wang, H. H., Carr, P. A., Mosberg, J. A., Rohland, N., Schultz, P. G., Jacobson, J. M., Rinehart, J., Church, G. M., & Isaacs, F. J. (2013). Genomically Recoded Organisms Expand Biological Functions. Science, 342(6156), 357. https://doi.org/10.1126/science.1241459

Ledoux, S., & Uhlenbeck, O. C. (2008, 2008/02/01/). [3′-32P]-labeling tRNA with nucleotidyltransferase for assaying aminoacylation and peptide bond formation. Methods, 44(2), 74–80. https://doi.org/10.1016/j.ymeth.2007.08.001

Lee, K. J., Kang, D., & Park, H.-S. (2019, May). Site-Specific Labeling of Proteins Using Unnatural Amino Acids. Molecules and Cells, 42(5), 386–396. https://doi.org/10.14348/molcells.2019.0078

Lerner, E., Cordes, T., Ingargiola, A., Alhadid, Y., Chung, S., Michalet, X., & Weiss, S. (2018, Jan). Toward dynamic structural biology: Two decades of single-molecule Forster resonance energy transfer. Science, 359(6373), 288-+, Article eaan1133. https://doi.org/10.1126/science.aan1133

Levi, M., Noel, J. K., & Whitford, P. C. (2019, Jun 1). Studying ribosome dynamics with simplified models. Methods, 162–163, 128-140. https://doi.org/10.1016/j.ymeth.2019.03.023

Liljeruhm, J., Wang, J., Kwiatkowski, M., Sabari, S., & Forster, A. C. (2019, 2019/02/15). Kinetics of d-Amino Acid Incorporation in Translation. ACS Chem Biol, 14(2), 204–213. https://doi.org/10.1021/acschembio.8b00952

Liu, C. C., & Schultz, P. G. (2010, 2010/06/07). Adding New Chemistries to the Genetic Code. Annual Review of Biochemistry, 79(1), 413–444. https://doi.org/10.1146/annurev.biochem.052308.105824

Liu, J., Hemphill, J., Samanta, S., Tsang, M., & Deiters, A. (2017). Genetic Code Expansion in Zebrafish Embryos and Its Application to Optical Control of Cell Signaling. Journal of the American Chemical Society, 139(27), 9100–9103. https://doi.org/10.1021/jacs.7b02145

Liu, T., Kaplan, A., Alexander, L., Yan, S., Wen, J.-D., Lancaster, L., Wickersham, C. E., Fredrick, K., Noller, H., Tinoco, I., Jr., & Bustamante, C. J. (2014, 2014/08/11). Direct measurement of the mechanical work during translocation by the ribosome. eLife, 3, e03406. https://doi.org/10.7554/eLife.03406

Louie, A., Masuda, E., Yoder, M., & Jurnak, F. (1984, Sep). Affinity purification of aminoacyl-tRNA. Analytical Biochemistry, 141(2), 402–408.

Maini, R., Chowdhury, S. R., Dedkova, L. M., Roy, B., Daskalova, S. M., Paul, R., Chen, S., & Hecht, S. M. (2015, 2015/06/16). Protein Synthesis with Ribosomes Selected for the Incorporation of β-Amino Acids. Biochemistry, 54(23), 3694–3706. https://doi.org/10.1021/acs.biochem.5b00389

Maini, R., Dedkova, L. M., Paul, R., Madathil, M. M., Chowdhury, S. R., Chen, S., & Hecht, S. M. (2015, 2015/09/09). Ribosome-Mediated Incorporation of Dipeptides and Dipeptide Analogues into Proteins in Vitro. Journal of the American Chemical Society, 137(35), 11206–11209. https://doi.org/10.1021/jacs.5b03135

Melnikov, S. V., Khabibullina, N. F., Mairhofer, E., Vargas-Rodriguez, O., Reynolds, N. M., Micura, R., Soll, D., & Polikanov, Y. S. (2019, Feb 28). Mechanistic insights into the slow peptide bond formation with D-amino acids in the ribosomal active site. Nucleic Acids Res, 47(4), 2089–2100. https://doi.org/10.1093/nar/gky1211

Melnikov, S. V., & Söll, D. (2019). Aminoacyl-tRNA Synthetases and tRNAs for an Expanded Genetic Code: What Makes them Orthogonal? International Journal of Molecular Sciences, 20(8). https://doi.org/10.3390/ijms20081929

Murakami, H., Ohta, A., Ashigai, H., & Suga, H. (2006, 2006/05/01). A highly flexible tRNA acylation method for non-natural polypeptide synthesis. Nature Methods, 3(5), 357–359. https://doi.org/10.1038/nmeth877

Neumann, H., Wang, K., Davis, L., Garcia-Alai, M., & Chin, J. W. (2010, 2010/03/01). Encoding multiple unnatural amino acids via evolution of a quadruplet-decoding ribosome. Nature, 464(7287), 441–444. https://doi.org/10.1038/nature08817

Obexer, R., Walport, L. J., & Suga, H. (2017, Jun). Exploring sequence space: harnessing chemical and biological diversity towards new peptide leads. Curr Opin Chem Biol, 38, 52–61. https://doi.org/10.1016/j.cbpa.2017.02.020

Ohtsuki, T., Yamamoto, H., Doi, Y., & Sisido, M. (2010, Aug). Use of EF-Tu mutants for determining and improving aminoacylation efficiency and for purifying aminoacyl tRNAs with non-natural amino acids. J Biochem, 148(2), 239–246. https://doi.org/10.1093/jb/mvq053

Ohuchi, M., Murakami, H., & Suga, H. (2007, 2007/10/01/). The flexizyme system: a highly flexible tRNA aminoacylation tool for the translation apparatus. Curr Opin Chem Biol, 11(5), 537–542. https://doi.org/10.1016/j.cbpa.2007.08.011

Orelle, C., Carlson, E. D., Szal, T., Florin, T., Jewett, M. C., & Mankin, A. S. (2015, 2015/08/01). Protein synthesis by ribosomes with tethered subunits. Nature, 524(7563), 119–124. https://doi.org/10.1038/nature14862

Park, H.-S., Hohn, M. J., Umehara, T., Guo, L.-T., Osborne, E. M., Benner, J., Noren, C. J., Rinehart, J., & Söll, D. (2011). Expanding the Genetic Code of Escherichia coli with Phosphoserine. Science, 333(6046), 1151. https://doi.org/10.1126/science.1207203

Pavlov, M. Y., & Ehrenberg, M. (1996, 1996/04/01/). Rate of Translation of Natural mRNAs in an Optimizedin VitroSystem. Archives of Biochemistry and Biophysics, 328(1), 9–16. https://doi.org/10.1006/abbi.1996.0136

Pavlov, M. Y., Watts, R. E., Tan, Z., Cornish, V. W., Ehrenberg, M., & Forster, A. C. (2009). Slow peptide bond formation by proline and other N-alkylamino acids in translation. Proceedings of the National Academy of Sciences, 106(1), 50. https://doi.org/10.1073/pnas.0809211106

Powers, T., & Noller, H. F. (1991, 1991/08/01). A functional pseudoknot in 16S ribosomal RNA. The EMBO Journal, 10(8), 2203–2214. https://doi.org/10.1002/j.1460-2075.1991.tb07756.x

Rackham, O., & Chin, J. W. (2005, 2005/08/01). A network of orthogonal ribosome·mRNA pairs. Nat Chem Biol, 1(3), 159–166. https://doi.org/10.1038/nchembio719

Ribeiro, S., Nock, S., & Sprinzl, M. (1995, Jul 1). Purification of aminoacyl-tRNA by affinity chromatography on immobilized Thermus thermophilus EF-Tu.GTP. Analytical Biochemistry, 228(2), 330–335.

Robertson, S. A., Ellman, J. A., & Schultz, P. G. (1991, 1991/03/01). A general and efficient route for chemical aminoacylation of transfer RNAs. Journal of the American Chemical Society, 113(7), 2722–2729. https://doi.org/10.1021/ja00007a055

Rogers, J. M., Kwon, S., Dawson, S. J., Mandal, P. K., Suga, H., & Huc, I. (2018, Apr). Ribosomal synthesis and folding of peptide-helical aromatic foldamer hybrids. Nat Chem, 10(4), 405–412. https://doi.org/10.1038/s41557-018-0007-x

Saleh, A. M., Wilding, K. M., Calve, S., Bundy, B. C., & Kinzer-Ursem, T. L. (2019, May 22). Non-canonical amino acid labeling in proteomics and biotechnology. Journal of Biological Engineering, 13, Article 43. https://doi.org/10.1186/s13036-019-0166-3

Schindelin, J., Arganda-Carreras, I., Frise, E., Kaynig, V., Longair, M., Pietzsch, T., Preibisch, S., Rueden, C., Saalfeld, S., Schmid, B., Tinevez, J.-Y., White, D. J., Hartenstein, V., Eliceiri, K., Tomancak, P., & Cardona, A. (2012, 2012/07/01). Fiji: an open-source platform for biological-image analysis. Nature Methods, 9(7), 676–682. https://doi.org/10.1038/nmeth.2019

Schmied, W. H., Tnimov, Z., Uttamapinant, C., Rae, C. D., Fried, S. D., & Chin, J. W. (2018, Dec). Controlling orthogonal ribosome subunit interactions enables evolution of new function. Nature, 564(7736), 444–448. https://doi.org/10.1038/s41586-018-0773-z

Stern, S., Moazed, D., & Noller, H. F. (1988). Structural analysis of RNA using chemical and enzymatic probing monitored by primer extension. Methods Enzymol, 164, 481–489. https://doi.org/10.1016/s0076-6879(88)64064-x

Takamoto, K., Chance, M. R., & Brenowitz, M. (2004, Aug 19). Semi-automated, single-band peak-fitting analysis of hydroxyl radical nucleic acid footprint autoradiograms for the quantitative analysis of transitions. Nucleic Acids Res, 32(15), E119. https://doi.org/10.1093/nar/gnh117

Tan, Z., Forster, A. C., Blacklow, S. C., & Cornish, V. W. (2004, 2004/10/01). Amino Acid Backbone Specificity of the Escherichia coli Translation Machinery. Journal of the American Chemical Society, 126(40), 12752–12753. https://doi.org/10.1021/ja0472174

Tharp, J. M., Ad, O., Amikura, K., Ward, F. R., Garcia, E. M., Cate, J. H. D., Schepartz, A., & Soll, D. (2020, Feb 17). Initiation of Protein Synthesis with Non-Canonical Amino Acids In Vivo. Angew Chem Int Ed Engl, 59(8), 3122–3126. https://doi.org/10.1002/anie.201914671

Tijerina, P., Mohr, S., & Russell, R. (2007). DMS footprinting of structured RNAs and RNA-protein complexes. Nat Protoc, 2(10), 2608–2623. https://doi.org/10.1038/nprot.2007.380

Tinoco, I., Jr., & Gonzalez, R. L., Jr. (2011, Jun 15). Biological mechanisms, one molecule at a time. Genes Dev, 25(12), 1205–1231.

Traut, R. R., & Monro, R. E. (1964, Oct). The puromycin reaction and its relation to protein synthesis. J Mol Biol, 10, 63–72. https://doi.org/10.1016/s0022-2836(64)80028-0

Tsiamantas, C., Rogers, J. M., & Suga, H. (2020, Apr 21). Initiating ribosomal peptide synthesis with exotic building blocks. Chem Commun (Camb), 56(31), 4265–4272. https://doi.org/10.1039/d0cc01291b

Tsurui, H., Kumazawa, Y., Sanokawa, R., Watanabe, Y., Kuroda, T., Wada, A., Watanabe, K., & Shirai, T. (1994, Aug 15). Batchwise purification of specific tRNAs by a solid-phase DNA probe. Anal Biochem, 221(1), 166–172. https://doi.org/10.1006/abio.1994.1393

van de Meent, J. W., Bronson, J. E., Wiggins, C. H., & Gonzalez, R. L., Jr. (2014, Mar 18). Empirical Bayes methods enable advanced population-level analyses of single-molecule FRET experiments. Biophys J, 106(6), 1327–1337. https://doi.org/10.1016/j.bpj.2013.12.055

Wagner, E. G. H., Jelenc, P. C., Ehrenberg, M., & Kurland, C. G. (1982, 1982/02/01). Rate of Elongation of Polyphenylalanine in vitro. European Journal of Biochemistry, 122(1), 193–197. https://doi.org/10.1111/j.1432-1033.1982.tb05866.x

Walker, A. S., Russ, W. P., Ranganathan, R., & Schepartz, A. (2020). RNA sectors and allosteric function within the ribosome. Proceedings of the National Academy of Sciences, 117(33), 19879. https://doi.org/10.1073/pnas.1909634117

Wang, L. (2017, Sep 25). Genetically encoding new bioreactivity. New Biotechnology, 38, 16–25. https://doi.org/10.1016/j.nbt.2016.10.003

Wang, L., Brock, A., Herberich, B., & Schultz, P. G. (2001, Apr 20). Expanding the genetic code of Escherichia coli. Science, 292(5516), 498–500.

Ward, F. R., Watson, Z. L., Ad, O., Schepartz, A., & Cate, J. H. D. (2019, 2019/11/12). Defects in the Assembly of Ribosomes Selected for β-Amino Acid Incorporation. Biochemistry, 58(45), 4494–4504. https://doi.org/10.1021/acs.biochem.9b00746

Weinger, J. S., Parnell, K. M., Dorner, S., Green, R., & Strobel, S. A. (2004, 2004/11/01). Substrate-assisted catalysis of peptide bond formation by the ribosome. Nature Structural & Molecular Biology, 11(11), 1101–1106. https://doi.org/10.1038/nsmb841

Wohlgemuth, I., Brenner, S., Beringer, M., & Rodnina, M. V. (2008, Nov 21). Modulation of the rate of peptidyl transfer on the ribosome by the nature of substrates. J Biol Chem, 283(47), 32229–32235. https://doi.org/10.1074/jbc.M805316200

Xu, Z., & Culver, G. M. (2009). Chemical probing of RNA and RNA/protein complexes. Methods Enzymol, 468, 147–165. https://doi.org/10.1016/s0076-6879(09)68008-3

